# Functional organization of multisensory integration network in children and youth with neurodevelopmental disorders predicts clinical sensory issues

**DOI:** 10.1101/2025.08.18.670938

**Authors:** Eun Jung Choi, Kathleen Lyons, Marlee Vandewouw, Evdokia Anagnostou, Paul D. Arnold, Muhammed Ayub, Jennifer Crosbie, Stelios Georgiades, Jessica Jones, Elizabeth Kelley, Azadeh Kushki, Jason P. Lerch, Russell J. Schachar, Bobby Stojanoski, Margot J. Taylor, Ryan A. Stevenson

**Author notes:** **Address correspondence to:** Ryan A. Stevenson, PhD Department of Psychology, University of Western Ontario 1137 Western Rd, London, Ontario N6G 1G7.

## Abstract

Differences in sensory processing in neurodevelopmental conditions, including Autism Spectrum Disorder (ASD) and Attention-deficit/Hyperactivity Disorder (ADHD), cascade into downstream clinical symptomatology. This includes differences in combining sensory information from multiple modalities into a unified percept, known as multisensory integration. Little is known about the functional organization of multisensory network (MSN) in these groups, its relation to clinical sensory issues, or its interaction with other higher-order cortical networks. We examined resting-state fMRI data from 417 participants in the Province of Ontario Neurodevelopmental Network (ASD=174, ADHD=130, Typical Development=113; Mean age=11.96±4.10). Timeseries data were extracted from the MSN and seven additional resting-state cortical networks (RSNs). Undirected and directed functional connectivity (FC) metrics were computed within the MSN and between the MSN and other RSNs. FC was compared across diagnoses and related to clinical sensory characteristics. The thalamus emerged as a hub region in both undirected and directed FC within the MSN and between the MSN-RSNs. Some diagnosis-related differences were observed, with increased MSN-RSN FC particularly in ADHD; however, associations with sensory characteristics were stronger in both undirected and directed FC within the MSN and between the MSN-RSNs, regardless of diagnosis. Converging evidence was seen in data-driven clusters based on FC metrics, which did not align with diagnosis, but instead mapped on to the overall level of sensory issues reported. That the data-driven clusters sorted not by diagnosis but by sensory characteristics suggests that these sensory characteristics and their underlying neurobiology are transdiagnostic in nature as opposed to specific to ASD or ADHD.

## Introduction

Sensory processing is a fundamental adaptive function that enables individuals to perceive, evaluate, and interpret sensory stimuli from the external environment, ultimately to generate an appropriate response. This process includes the detection and encoding of sensory inputs from peripheral receptors (e.g., visual and auditory receptors), and subsequent integration of sensory inputs across modalities.^1^ Given the dynamic and complex nature of real-world environments, multisensory integration allows the brain to combine information from different sensory modalities for more efficient and accurate perception and response. Neuroimaging evidence demonstrates that multisensory integration occurs not only in higher-order associative regions but also in early sensory cortices, suggesting that the sensory systems inherently accommodate environmental demands by processing multisensory information concurrently at multiple hierarchical levels. ^2,3^ When this system is disrupted, the formation of accurate perceptual representations and appropriate behavioural responses can be compromised.^4^

Multisensory integration also serves as a fundamental building block for higher-order cognitive functions, including attention, memory, learning, and language.^5–7^ For instance, early communicative skills such as attending to caregivers in social situations and object naming require integrating multisensory information.^8^ Disruptions in multisensory integration have cascading effects on higher-order cognition, suggesting sensory difficulties play a precursor role in various cognitive and social difficulties observed in neurodevelopmental conditions.^9–13^ However, the underlying neurobiology of how the multisensory network interacts with other higher-order cortical networks remains poorly understood. Furthermore, it is unclear to what extent these interactions are similar or distinct across different neurodevelopmental disorders.

Sensory difficulties are one of the core diagnostic profiles in Autism Spectrum Disorder (ASD),^14^ with up to 95% of autistic individuals experiencing hyper-and hypo-sensitivity across multiple sensory modalities, as well as sensory-seeking behaviours.^15–17^ Atypical multisensory integration has been reported perceptually through behavioural studies, and is associated with altered neural activation, disrupted functional connectivity (FC), and an imbalance in excitatory/inhibitory signaling across sensory and multisensory processing regions.^13^ Neuroimaging studies have revealed large-scale functional alterations, including reduced activation in the superior temporal sulcus, one of the multisensory hub regions,^18,19^ and weaker long-range connectivity between sensory cortices and higher-order cognitive regions in the default mode network, relative to neurotypical patterns.^13,20^ These findings suggest that atypical functional organization between multisensory and higher-order cortical regions may underlie complex clinical manifestations in this condition, spanning both lower-level sensory processing differences and higher-level social and cognitive impairments.^21^ Further, these differences in multisensory integration have been directly linked to core clinical features of ASD such as differences in social communication and repetitive behaviours.^10,22–24^

Atypical sensory processing is not exclusive to ASD but also observed in other neurodevelopmental disorders, such as Attention-Deficit/Hyperacitivity Disorder (ADHD). Children with ADHD exhibit sensory difficulties compared to their typically developing peers.^25,26^ Sensory challenges in ADHD have been reported across sensory modalities,^26–29^ and are present in up to 69% of children with ADHD.^30^ Moreover, similar sensory phenotypes and difficulties have been reported in both children with ASD and ADHD, suggesting transdiagnostic features in sensory processing atypicalities.^25^ Children with ADHD also exhibit difficulties in multisensory integration, showing lower susceptibility to the McGurk effect,^31–33^ and diminished neural responses during the multisensory integration tasks.^34^ These behavioural differences align with neuroimaging findings indicating functional alterations in unimodal and multisensory sensory areas^32^ as well as maturational structural delay in fronto-temporal connectivity, between Heschl’s gyrus and auditory parabelt regions, suggesting disruptions in auditory processing and integration.^31^

While multisensory integration difficulties in both ASD and ADHD are well documented, the shared and distinct neurobiological mechanisms underlying these challenges is less understood. Some common findings of specific neural substrates, such as alterations in the superior temporal cortices, have implicated the underlying mechanisms across the conditions since they play a critical role in audiovisual integration.^35–38^ However, few studies have examined how multisensory regions interact and are functionally organized with higher-order network at a large-scale level. A broader, network-level perspective is needed to understand how multisensory integration difficulties relate to higher-order cognitive differences in ASD and ADHD.

Traditional FC measures how two brain regions co-activate over time, capturing undirected synchronous activity. However, evidence for neural mechanisms on multisensory integration (e.g., phase coupling and amplitude modulation) suggests temporal dynamics play a critical role in large-scale network organization.^1^ Understanding the directed influences between multisensory regions and other cortical areas is essential for understanding information flow.^39^ One approach for assessing directed associations is Granger causality, which determines whether neural activity in one brain region at a designated time point can predict the activity of another region at a subsequent time point.^40,41^ This method leverages temporal dependencies to provide insights into potential causal influences between regions. Together, undirected and directed FC offer complementary perspectives on the brain’s functional architecture, from co-activation patterns to directional information transfer.

The present study examined the functional organization of key multisensory integration regions in children with ASD and ADHD, and typical development. Both undirected and directed FC were explored within the multisensory network (MSN) as well as in the associations with other resting-state networks (RSNs). We investigated diagnostic differences in network organization, related FC patterns to clinical sensory issues, and applied a data-driven clustering approach based solely on the functional organizational features, regardless of diagnoses, to determine whether the resulting clusters aligned with clinical diagnoses and/or sensory processing abilities.

## Methods

### Participants

Data were from the Province of Ontario Neurodevelopmental Disorder network (POND), a research collaborative group in Ontario, Canada. The brain imaging data included that children and youth who had no clinical diagnoses (TD) or had a diagnosis of ASD or ADHD. POND participants with ASD, ADHD, or TD who completed both resting-state fMRI and Short Sensory Profile (SSP)^42^ were included, with a total N of 505 (ASD=226, ADHD=150, TD=129).

### Data acquisition

The resting-state fMRI data were collected in three different protocols, 3T TimTrio while children and youth watching a movie (N=37, sequence=EPI, TR=2340ms, TE=30ms, FA=70, FOV(mm)=224×224×140, voxel size=3.5mm, scan time=5 min), 3T TimTrio while children and youth watching a naturalistic movie paradigm called Inscapes^43^ (N=94, sequence=EPI, TR=2340ms, TE=30ms, FA=70, FOV(mm)=224×224×140, voxel size=3.5mm, scan time=5 min), and 3T Prisma FIT while children and youth watching Inscapes (N=286, sequence=EPI, TR=1500ms, TE=30ms, FA=70, FOV(mm)=224×224×150, voxel size=3mm, scan time=5 min).

### Data processing

Anatomical and functional fMRI data were preprocessed using fMRIPrep^44^. Anatomical preprocessing included intensity non-uniformity correction, skull-stripping, spatial normalization to a paediatric template, and segmentation into cerebrospinal fluid, white matter, and gray matter.

Functional preprocessing involved slice-time correction, motion correction, co-registration to the T1-weighted image, and exclusion of frames exceeding motion thresholds. Participants with more than 1/3 of their frames exceeding the recommended threshold (FD:0.5mm, DVARS: 1.5) were excluded from all subsequent analyses, resulting in 417 participants remaining (ASD=174, ADHD=130, TD=113). Nuisance regressors (e.g., motion parameters, CSF, WM) were removed, and temporal filtering was applied (Supplementary information for more details).

### Constructing FC maps

The MSN was defined based on a recent study that identified brain regions involved in multisensory integration across multiple sensory modalities (auditory, visual, tactile, gustatory, and olfactory) using an Activation Likelihood Estimation meta-analysis.^45^ The key regions include the superior temporal gyri/sulci, middle temporal gyri, thalamus, right insula, and left inferior frontal gyrus, identified through interaction effects for multisensory stimuli (e.g., superadditive, conjunction contrasts, or maximum/mean criteria),^46^ suggesting FC within the MSN. The RSNs were defined using Yeo’s seven RSNs, including fronto-parietal, default-mode, dorsal attention, limbic, ventral attention, somatomotor, and visual networks.^47^ These networks were identified through a large-scale FC analysis of resting-state fMRI data, which clustered cortical regions into seven distinct networks based on their correlated activity patterns. Timeseries data were extracted from five masks in the MSN^45^ and fifty-one regional masks in Yeo’s seven RSNs^47^(Figure 1).

**Figure 1.**
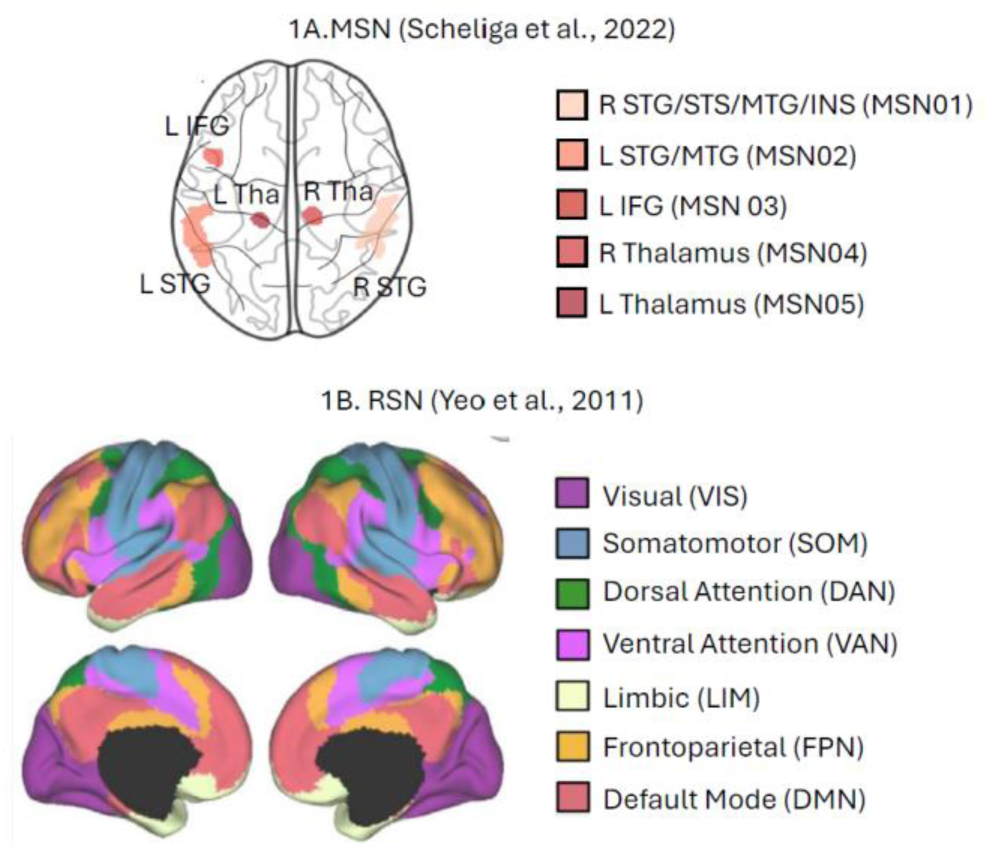
Definition of MSN and RSN. MSN, Multisensory Network; RSN, Resting-state network; R, Right hemisphere; L, Left hemisphere; STG, Superior temporal gyrus; STS, Superior temporal sulcus; MTG, Middle temporal gyrus; INS, Insula; IFG, Inferior frontal gyrus; Tha, Thalamus

The within-MSN FC was defined in 5 by 5 ROI pairings, resulting in 10 unique undirected FC pairings and 20 unique directed FC pairings (Supplementary Table 1). The between MSN-RSN FC was defined in 5 by 51 ROI pairings, resulting in 255 unique undirected FC and 510 unique directed FC pairs (Supplementary Table 2). The undirected connectivity was computed in pairwise Pearson’s correlation coefficients between the regions and converted to a z-value using Fisher’s r-to-z transform to better approximate a normal distribution to represent within-MSN and between MSN-RSN connectivity. The directed connectivity was calculated in Granger causality using MVGC toolbox.^41^ To compute Granger causality, we preprocessed the timeseries data by first applying linear detrending to remove long-term trends, followed by z-score normalization to ensure comparability across regions of interest (ROIs) and participants. Stationarity was assessed using the Augmented Dickey-Fuller test, and non-stationary timeseries data were transformed via first-order differencing to meet the assumptions of Granger causality analysis. Missing data points were addressed through linear interpolation, ensuring complete timeseries data. Lag selection was performed using Vector Autoregression models, with the maximum lag set to 10% of the timeseries length for each participant to account for variability in dataset size. Optimal lags were identified using Bayesian Information Criterion (BIC), selecting the lag that minimized each criterion to balance model fit and complexity, resulting in BIC=1. To account for the effects of different scanners, corrections on the constructed FC maps were made for the different acquisition scanners using ComBat harmonization^48^, during which no biological variables were included as fixed effects, and the default parametric prior method was used in the empirical Bayes procedure.

### Sensory processing abilities

Participants’ sensory processing abilities were assessed using the Short Sensory Profile (SSP)^42^, a widely validated parent-report questionnaire. This tool evaluates behaviours related to atypical sensory responses in everyday life across seven domains: tactile sensitivity, taste/smell sensitivity, movement sensitivity, under-responsiveness and sensation-seeking, auditory filtering, low energy/weakness, and visual/auditory sensitivity (Supplementary information).

### Analysis

Both undirected and directed functional connectivity (FC) within-MSN and between-MSN-RSN were examined in three steps: (1) diagnostic differences while controlling for age and sex, (2) associations with sensory processing abilities while controlling for age and sex, and (3) data-driven FC clusters and their associations with diagnoses and sensory processing abilities. Diagnostic differences were tested using analysis of covariance (ANCOVA), with age and sex included as covariates. Regression analyses were conducted to examine the associations between FC and sensory processing abilities, also controlling for age and sex. For both ANCOVA and regression analyses, 5000 permutations were applied. Specifically, group labels (for ANCOVA) or sensory scores (for regression) were shuffled across participants while keeping covariates fixed. For each permutation, the test statistic was recalculated to create a null distribution, and p-values were assigned based on the proportion of permuted test statistics that exceeded the observed statistic. Results were considered significant at p < 0.05. The effect sizes were reported in either partial eta squared or semi-partial *R*^2^. For data-driven analyses, elbow and silhouette methods were used to determine the optimal number of clusters, followed by k-means and k-medoids, depending on data normality.^49,50^ Participants’ diagnoses and sensory processing abilities were used to characterize group membership in each cluster. For each pairwise comparison, False Discovery Rate (FDR) corrections were performed using Holm-Bonferroni^51^, q < 0.05, to correct for comparisons across multiple networks.

## Results

### Participants

The sample included in the final analysis was 417 (ASD=174, ADHD=130, TD=113; Mean age=11.96±4.10 years; 32% female). Demographic information is presented in Table 1. The three groups did not differ in age; however, the ASD and ADHD groups included a higher proportion of male participants (χ²=18.3, p<0.001).^52^ The ASD group exhibited the greatest difficulties across all sensory domains, while the ADHD group also showed significant sensory processing challenges (tactile, taste/smell, movement, under-responsiveness/sensation-seeking, auditory filtering, low energy/weakness, and visual/auditory). Notably, the ADHD group did not differ from the ASD group in the under-responsive/seeks sensation and auditory filtering domains.

**Table 1.**
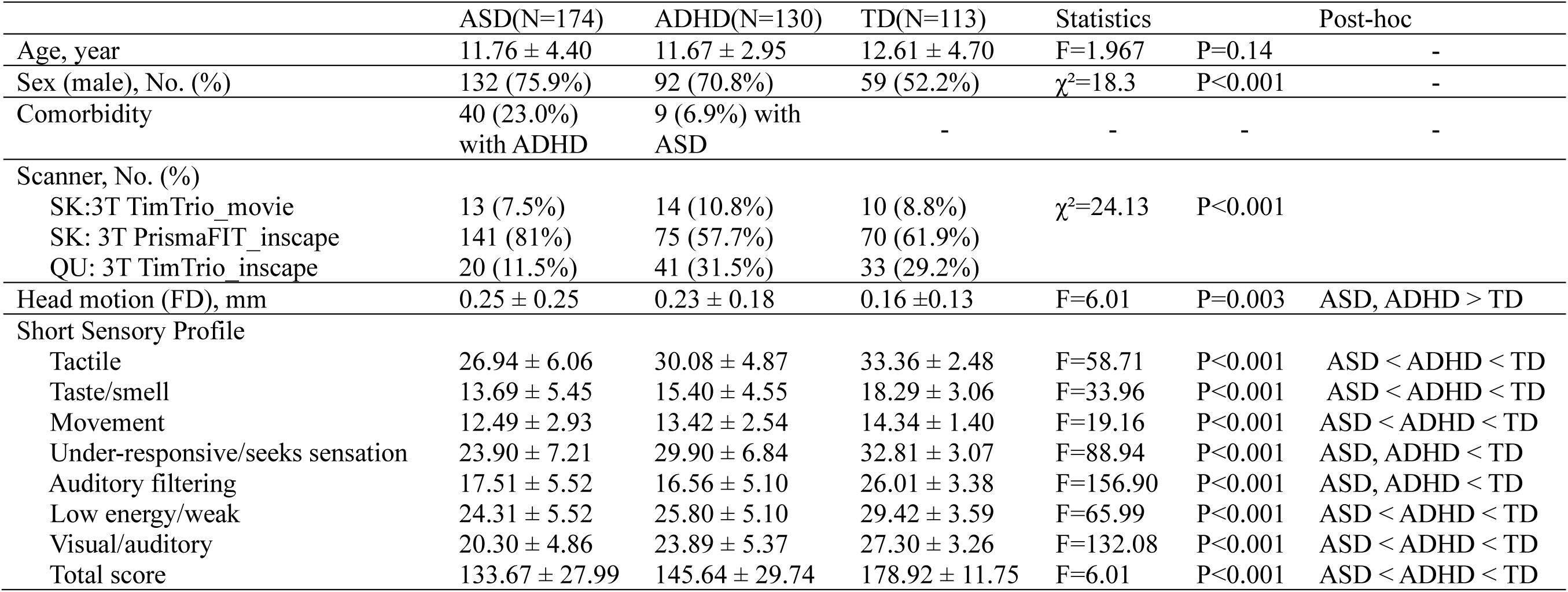
Descriptive statistics for participants demographics

|  | ASD(N=174) | ADHD(N=130) | TD(N=113) | Statistics |  | Post-hoc |
| --- | --- | --- | --- | --- | --- | --- |
| Age, year | 11.76 ± 4.40 | 11.67 ± 2.95 | 12.61 ± 4.70 | F=1.967 | P=0.14 | - |
| Sex (male), No. (%) | 132 (75.9%) | 92 (70.8%) | 59 (52.2%) | $\chi^2=18.3$ | P<0.001 | - |
| Comorbidity | 40 (23.0%)<br>with ADHD | 9 (6.9%) with<br>ASD | - | - | - | - |
| Scanner, No. (%) |  |  |  |  |  |  |
| SK:3T TimTrio_movie | 13 (7.5%) | 14 (10.8%) | 10 (8.8%) | $\chi^2=24.13$ | P<0.001 | |
| SK: 3T PrismaFIT_inscape | 141 (81%) | 75 (57.7%) | 70 (61.9%) |  |  |  |
| QU: 3T TimTrio_inscape | 20 (11.5%) | 41 (31.5%) | 33 (29.2%) |  |  |  |
| Head motion (FD), mm | 0.25 ± 0.25 | 0.23 ± 0.18 | 0.16 ± 0.13 | F=6.01 | P=0.003 | ASD, ADHD > TD |
| Short Sensory Profile |  |  |  |  |  |  |
| Tactile | 26.94 ± 6.06 | 30.08 ± 4.87 | 33.36 ± 2.48 | F=58.71 | P<0.001 | ASD < ADHD < TD |
| Taste/smell | 13.69 ± 5.45 | 15.40 ± 4.55 | 18.29 ± 3.06 | F=33.96 | P<0.001 | ASD < ADHD < TD |
| Movement | 12.49 ± 2.93 | 13.42 ± 2.54 | 14.34 ± 1.40 | F=19.16 | P<0.001 | ASD < ADHD < TD |
| Under-responsive/seeks sensation | 23.90 ± 7.21 | 29.90 ± 6.84 | 32.81 ± 3.07 | F=88.94 | P<0.001 | ASD, ADHD < TD |
| Auditory filtering | 17.51 ± 5.52 | 16.56 ± 5.10 | 26.01 ± 3.38 | F=156.90 | P<0.001 | ASD, ADHD < TD |
| Low energy/weak | 24.31 ± 5.52 | 25.80 ± 5.10 | 29.42 ± 3.59 | F=65.99 | P<0.001 | ASD < ADHD < TD |
| Visual/auditory | 20.30 ± 4.86 | 23.89 ± 5.37 | 27.30 ± 3.26 | F=132.08 | P<0.001 | ASD < ADHD < TD |
| Total score | 133.67 ± 27.99 | 145.64 ± 29.74 | 178.92 ± 11.75 | F=6.01 | P<0.001 | ASD < ADHD < TD |

### Undirected FC

The undirected FC maps within and between MSN and RSNs in each group are presented in Supplementary Figure 1. Within the MSN, a general patten of stronger connectivity was observed between inter-hemispheric homologues, such as bilateral superior temporal gyri and thalami in all participants. In the between MSN-RSN FC, the higher-order cortical regions in the MSN exhibited stronger connections, not only with sensory-related networks (somatomotor and visual networks) but also with dorsal and ventral attention networks. In contrast, the bilateral thalami in the MSN showed stronger connections with sensory-related networks compared to other higher-order cortical regions in the RSNs.

#### Diagnostic differences

All three groups demonstrated similar patterns of connectivity matrices within the MSN (**Figure 2**A). In the within-MSN FC, the main effect of diagnosis was only significant for the FC between the left STG and the right thalamus (F=3.10, p=0.05) with a small effect size (partial η²=0.01), while controlling for age and sex (Supplementary Table 3). In the post-hoc analyses, although both ASD and ADHD groups showed increased FC compared to the TD group, only ADHD group was significantly different (t=2.82, p=0.02) (**Figure 2**B).

**Figure 2.**
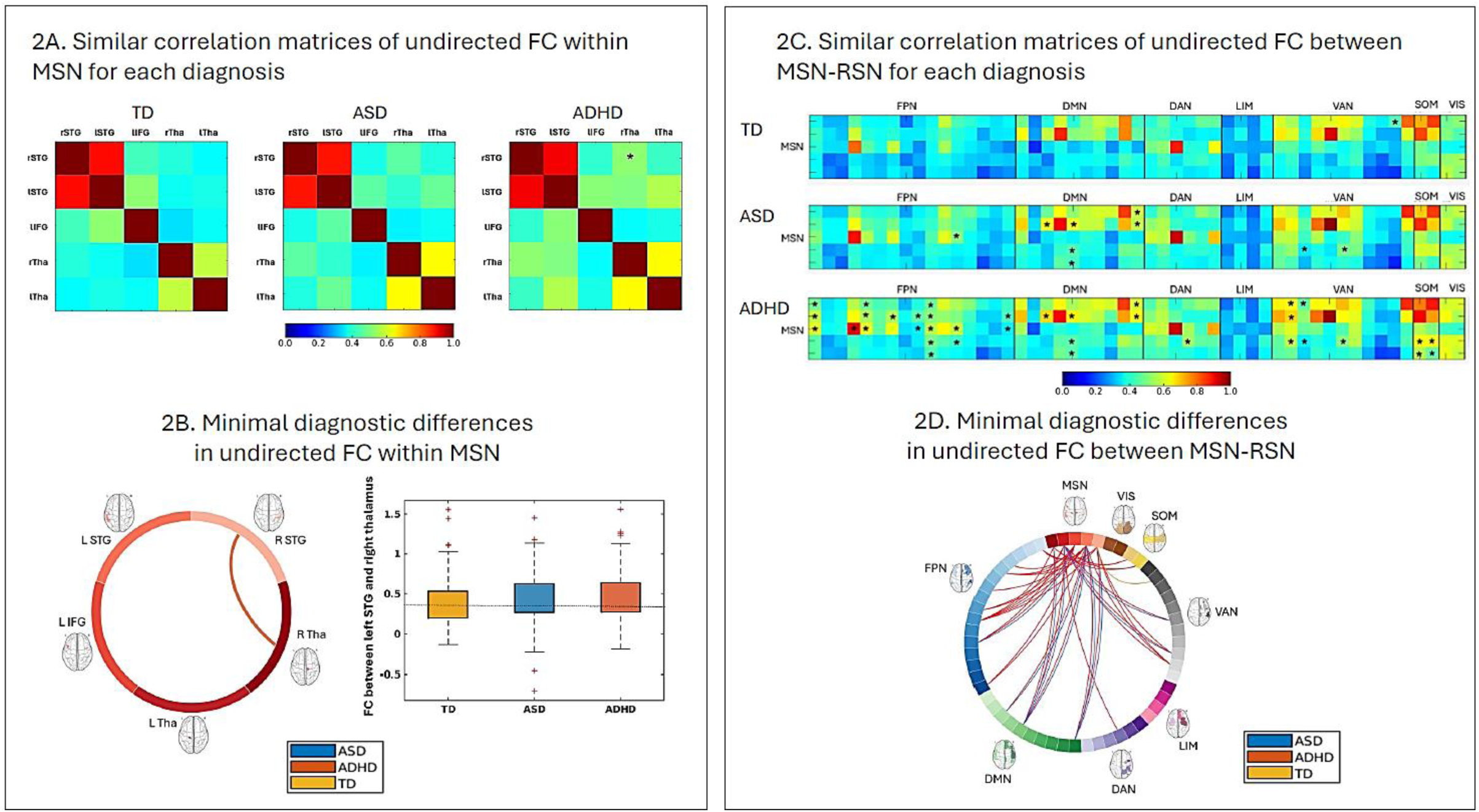
Diagnostic differences in undirected FC within the MSN and between the MSN and RSNs. **2A**. All three groups exhibited similar patterns of FC within the MSN, **2B**. Minimal diagnostic difference was found only in the FC between rSTG and rTha, with post-hoc analyses revealing significantly greater FC in ADHD group compared to the other groups (red line indicates greater FC in the ADHD group), **2C**. Across groups, overall FC patterns between the MSN and RSNs were largely similar, **2D**. Minimal diagnostic differences in FC between MSN and RSNs were observed mostly in the ADHD group. Red lines indicate connections with greater FC in the ADHD group, blue lines indicate greater FC in the ASD group, and the yellow line indicates greater FC in the TD group. rSTG, Right Superior Temporal Gyrus; lSTG, Left Superior Temporal Gyrus; lIFG, Left Inferior Frontal Gyrus; rTha, Right Thalamus; lTha, Left Thalamus; FPN, Fronto-parietal network; DMN, Default-mode network; DAN, Dorsal attention network; LIM, Limbic network; VAN, Ventral attention network; SOM, Somatomotor network; VIS, Visual network; *statistically significance at p<0.05 in diagnostic differences after controlling age and sex

Correlation matrices of between MSN-RSN FC for each diagnosis also presented similar patterns overall (**Figure 2**C). In the between MSN-RSN FC, significant main effects of diagnosis were observed in 37 connections among the 255 regional FCs across RSNs, excluding the LIM and VIS networks, while controlling for age and sex (Supplementary Table 4). In the FPN, diagnostic differences were found across all five MSN regions, with small to medium effect sizes (partialη²=0.01–0.03). In the DMN, these differences involved the bilateral STG and thalamus, with small effect sizes (partialη²=0.01–0.02). In the DAN, diagnostic differences were limited to connections with the right thalamus (partialη²=0.03). The VAN showed differences in FC with the bilateral STG and right thalamus, also with small effect sizes (partialη²=0.01–0.02). Lastly, in the SOM network, differences were observed with the bilateral thalamus (partialη²=0.02). Post-hoc analyses revealed that the ADHD group consistently exhibited increased FC in nearly all identified connections. While the ASD group also showed overall increased FC, the DMN was particularly notable for its heightened FC in ASD (**Figure 2**D).

#### Associations with sensory processing abilities

A substantial number of FC measures were associated with sensory processing abilities in both within-MSN FC and between MSN-RSN FC, with overall greater effect sizes compared to diagnostic differences. In the within-MSN FC, negative associations were observed between the bilateral thalamus and other MSN sub-regions and SSP total scores, after controlling for age and sex.

Specifically, increased FC between the right thalamus and the right STG (β=-0.001, p=0.01), left STG (β=-0.001, p=0.003), and left IFG (β=-0.001, p=0.004) was linked to poorer sensory processing abilities. Similarly, increased FC between the left thalamus and the right STG (β=-0.001, p=0.003), left STG (β=-0.002, p=0.0002), and left IFG (β=-0.001, p=0.02) was also associated with poorer sensory processing abilities. These associations demonstrated small to medium effect sizes (partialR²=0.03–0.06), emphasizing the role of thalamic connections in sensory processing (Supplementary Table 5).

In the analysis of between MSN-RSN FC, significant negative associations were observed between SSP total scores and FC across all MSN-RSN connections, indicating that increased MSN-RSN FC was associated with poorer sensory processing abilities. Among the RSNs, the FPN demonstrated significant associations with all five MSN sub-regions, with partial R² values ranging from 0.02 to 0.06, reflecting small to medium effect sizes. Similarly, the DMN exhibited significant associations with all MSN sub-regions, with partial R² values ranging from 0.02 to 0.07, including a strong effect in the FC between the right temporal cortex in the DMN and the right STG in the MSN (partialR²=0.07). The DAN consistently showed associations across its connections with MSN sub-regions, with partial R² values ranging from 0.02 to 0.03, representing small effect sizes. Likewise, the VAN revealed associations with small to medium effect sizes, with partial R² values between 0.01 and 0.05. Associations with the SOM and VIS networks were also observed, with partial R² values ranging from 0.02 to 0.03 for SOM and 0.01 to 0.04 for VIS, reflecting small to medium effect sizes. In contrast, the LIM network demonstrated a more limited association, with only the left thalamus in the MSN significantly associated with SSP scores (partialR²=0.02) (Supplementary Table 6).

#### Data-driven clusters and their associations with diagnosis and sensory processing abilities

In the within-MSN FC analysis, the elbow method identified three optimal clusters (Supplementary Figure 2A), each exhibiting distinct FC patterns (**Figure 3**A). The overall FC strength across the five MSN sub-regions increased in the order of Cluster2 (N=191) < Cluster1 (N=183) < Cluster3 (N=43).

**Figure 3.**
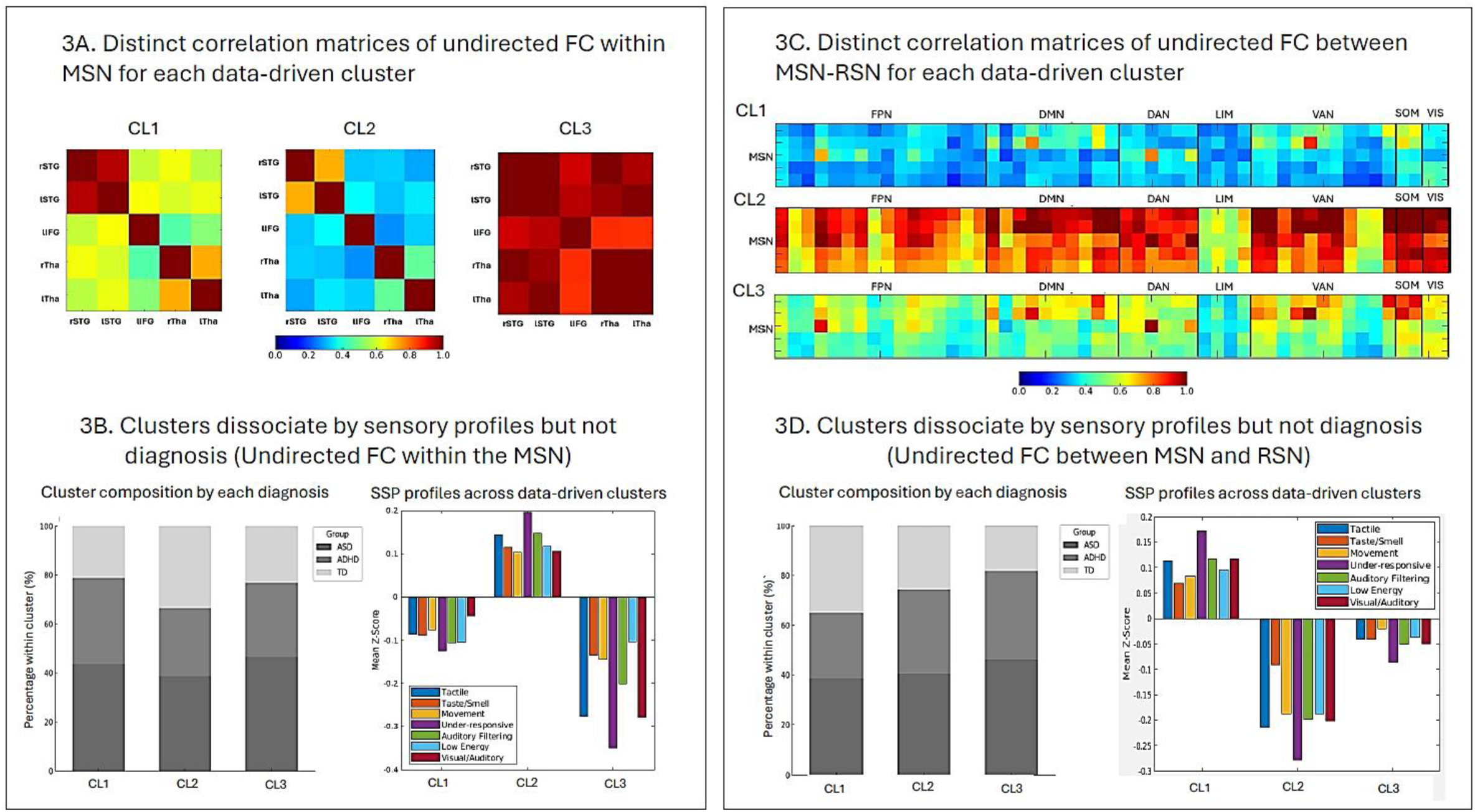
Data-driven clusters based on undirected FC withn the MSN and between the MSN and RSNs. **3A**. Three data-driven clusters exhibited distinct patterns of FC within the MSN, **3B**. When normalized by cluster size (%), all three clusters included participants from each diagnostic group, but demonstrated distinct sensory profile features: Cluster 1 (mild sensory issues across all domains), Cluster 2 (sensory adaptive), and Cluster 3 (generalized sensory issues), **3C**. Three data-driven clusters also showed distinct FC patterns between the MSN and RSNs, **3D**. When normalized by cluster size (%), each cluster again included all diagnostic groups, but demonstrated distinct sensory profiles: Cluster 1 (sensory adaptive), Cluster 2 (generalized sensory issues), and Cluster 3 (mild sensory issues across all domains) rSTG, Right Superior Temporal Gyrus; lSTG, Left Superior Temporal Gyrus; lIFG, Left Inferior Frontal Gyrus; rTha, Right Thalamus; lTha, Left Thalamus; FPN, Fronto-parietal network; DMN, Default-mode network; DAN, Dorsal attention network; LIM, Limbic network; VAN, Ventral attention network; SOM, Somatomotor network; VIS, Visual network

Despite these differences, the clusters did not significantly differ by age (F=2.29, p=0.10), sex (χ²=3.31, p=0.19), or scanner (χ²=3.24, p=0.52). Within each diagnostic group, the three clusters were proportionally distributed (Cluster1: Cluster2: Cluster3): TD=34.51%, 56.64%, 8.85%; ASD=45.98%, 42.53%, 11.49%; ADHD=49.23%, 40.77%, 10% (χ²=7.79, p=0.09). Notably, the sensory profiles provided clearer distinctions between clusters than diagnostic group distributions (**Figure 3**B).

Similarly, the between MSN-RSN FC analysis identified three optimal clusters (Supplementary Figure 2B), with distinct FC patterns (**Figure 3**C). The FC strength across MSN-RSN connections increased in the order of Cluster1 (N=189) < Cluster3 (N=160) < Cluster2 (N=68). The clusters did not significantly differ by age (F=1.39, p=0.25) or scanner (χ²=1.31, p=0.86), but sex differences were significant (χ²=7.89, p=0.02), with females more likely to belong to Cluster 1. Within each diagnostic group, the three clusters were proportionally distributed (Cluster1: Cluster2: Cluster3): TD=58.41%, 17.24%, 40.81%; ASD=41.95%,17.24%, 40.81%; ADHD=38.46%, 19.23%, 42.31% (χ²=4.12, p=0.39). As with the within-MSN analysis, sensory profiles were more distinct across clusters than diagnostic distributions (**Figure 3**D).

### Directed FC

The whole directed functional connectivity maps within the MSN and between MSN and RSNs in each group are presented in Supplementary Figure 3. Notably, the role of the thalamus was prominent in directed connectivity, with the right thalamus exhibiting greater directed FC to other multisensory regions within the MSN as well as to nearly all other regions in the RSNs.

#### Diagnostic differences

In the within-MSN FC, overall FC patterns were similar across diagnostic groups and the directions from right thalamus to other sub-regions in MSN were greater across all three groups (**Figure 4**A). The diagnostic differences were not observed in any of directed FC while controlling for age and sex (**Figure 4**B and Supplementary Table 7).

**Figure 4.**
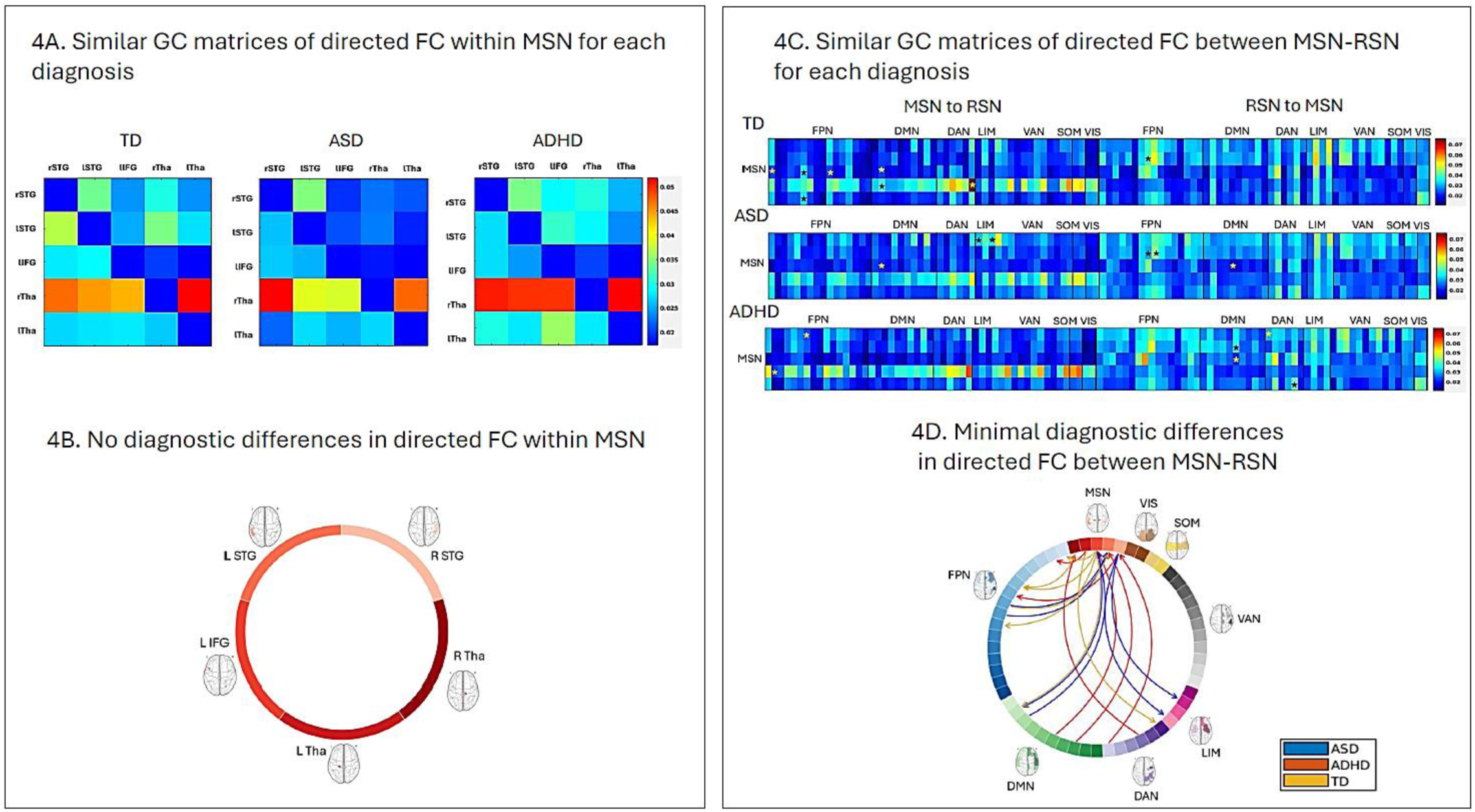
Diagnostic differences in directed FC within the MSN and between the MSN and RSNs. **4A**. All three groups exhibited similar patterns of directed FC within the MSN, with a shared trend of greater directionality from the right thalamus to other MSN subregions, **4B**. No significant group differences were observed in directed FC within the MSN, **4C.** Similar patterns were also observed between the MSN and RSNs across all groups, again showing a common trend of greater directionality from the right thalamus to other RSN subregions, **4D**. Minimal diagnostic differences in directed FC between the MSN and RSNs were identified. Each color represents a diagnostic group, and arrowheads indicate the direction of connectivity between the regions. Red lines indicate greater directional FC in the ADHD group, blue lines indicate greater directional FC in the ASD group, and yellow lines indicate greater directional FC in the TD group GC, Granger Causality; rSTG, Right Superior Temporal Gyrus; lSTG, Left Superior Temporal Gyrus; lIFG, Left Inferior Frontal Gyrus; rTha, Right Thalamus; lTha, Left Thalamus; FPN, Fronto-parietal network; DMN, Default-mode network; DAN, Dorsal attention network; LIM, Limbic network; VAN, Ventral attention network; SOM, Somatomotor network; VIS, Visual network; *statistically significance at p<0.05 in diagnostic differences after controlling age and sex

In the between MSN-RSN FC, overall FC patterns were also similar across diagnostic groups, with a shared trend of greater directionality originating from right thalamus toward other RSN subregions (**Figure 4**C). Significant main effects of diagnosis were observed in both MSN-to-RSN and RSN-to-MSN directed connections while controlling for age and sex (**Figure 4**D and Supplementary Table 8).

MSN-to-RSN FC: The TD group showed greater directed FC compared to the diagnostic groups in the directions from left IFG to FPN (partial η²=0.01–0.02) and DMN (partialη²=0.02), as well as from right thalamus to DAN (partialη²=0.02). However, the directions from the right STG to FPN were reversely increased in ADHD compared to other groups (partialη²=0.02). Notably, in the LIM network, the ASD group exhibited increased directed FC from the rSTG to LIM compared to both the TD and ADHD groups (partialη²=0.03).

RSN-to-MSN FC: In contrast to the directed FC from MSN to RSN, there was an overall increase in directed FC from RSN to MSN in both the ASD and ADHD groups. Specifically, the ASD group exhibited greater directed FC from FPN to MSN (left STG) compared to the ADHD group (partialη²=0.01-0.02). Additionally, increased directed FC from DMN to MSN (left IFG) was observed in the ASD group compared to both the ADHD and TD groups (partialη²=0.01-0.02). The ADHD group demonstrated enhanced directed FC from DMN to MSN (left STG) relative to the TD group (partialη²=0.01-0.03), as well as from DAN (right STG and thalamus) to MSN compared to both ASD and TD groups (partialη²=0.01-0.02].

#### Associations with sensory processing abilities

In the within-MSN FC, only the directed FC from the left STG was significantly associated with sensory processing abilities. Specifically, connections to the left IFG (β=-0.0001, p=0.05) and the right thalamus (β=0.0002, p=0.005) were observed, with small effect sizes (partialR²=0.01-0.02) (Supplementary Table 9).

In the between MSN-RSN FC, the sensory processing abilities were associated with connections between MSN and higher-order RSNs such as FPN, DMN, DAN and VAN in both directions.

MSN-to-RSN FC: Greater directed FC from STG and IFG in the MSN to the FPN (partialR²=0.02) and the VAN (partialR²=0.01-0.02) was positively associated with sensory processing abilities.

Conversely, greater directed FC from thalamus to the FPN (partial R² =0.02) and the DAN (partial R²=0.01-0.02) was negatively associated with sensory processing abilities (Supplementary Table 10).

RSN-to-MSN FC: Opposite patterns were observed in the RSN-to-MSN FC. Greater directed FC from the FPN (partialR²=0.02) and DMN (partialR²=0.02-0.04) to the STG and/or IFG was negatively associated with sensory processing abilities. However, greater directed FC from the FPN (partialR²=0.02) and DMN (partialR²=0.02) to the thalamus was positively associated with sensory processing abilities. Lastly, the connections between DAN and left thalamus demonstrated negative associations with sensory processing abilities in all directions. Similarly, greater directed FC from the Lim to the right thalamus was negatively associated with sensory processing abilities.

#### Data-driven clusters and their associations with diagnosis and sensory processing abilities

In the within-MSN FC analysis, the silhouette method identified two optimal clusters (Supplementary Figure 4A), each exhibiting distinct FC patterns (**Figure 5**A). The overall FC strength across the five MSN sub-regions increased in the order of Cluster1 (N=326) < Cluster2 (N=91), particularly increased directed FC from right thalamus to STG/IFG (Figure 7A). Despite these differences, the clusters did not significantly differ by age (t=-0.62, p=0.53), sex (χ²=0.20, p=0.66), or scanner (χ²=0.25, p=0.88). Within each diagnostic group, the two clusters were proportionally distributed (Cluster1: Cluster2): TD=76.99%, 23.01%; ASD=77.01%, 22.99%; ADHD=80.77%, 19.23% (χ²=0.74, p=0.67). Notably, sensory profiles were more distinct across clusters than diagnostic distributions (**Figure 5**B).

**Figure 5.**
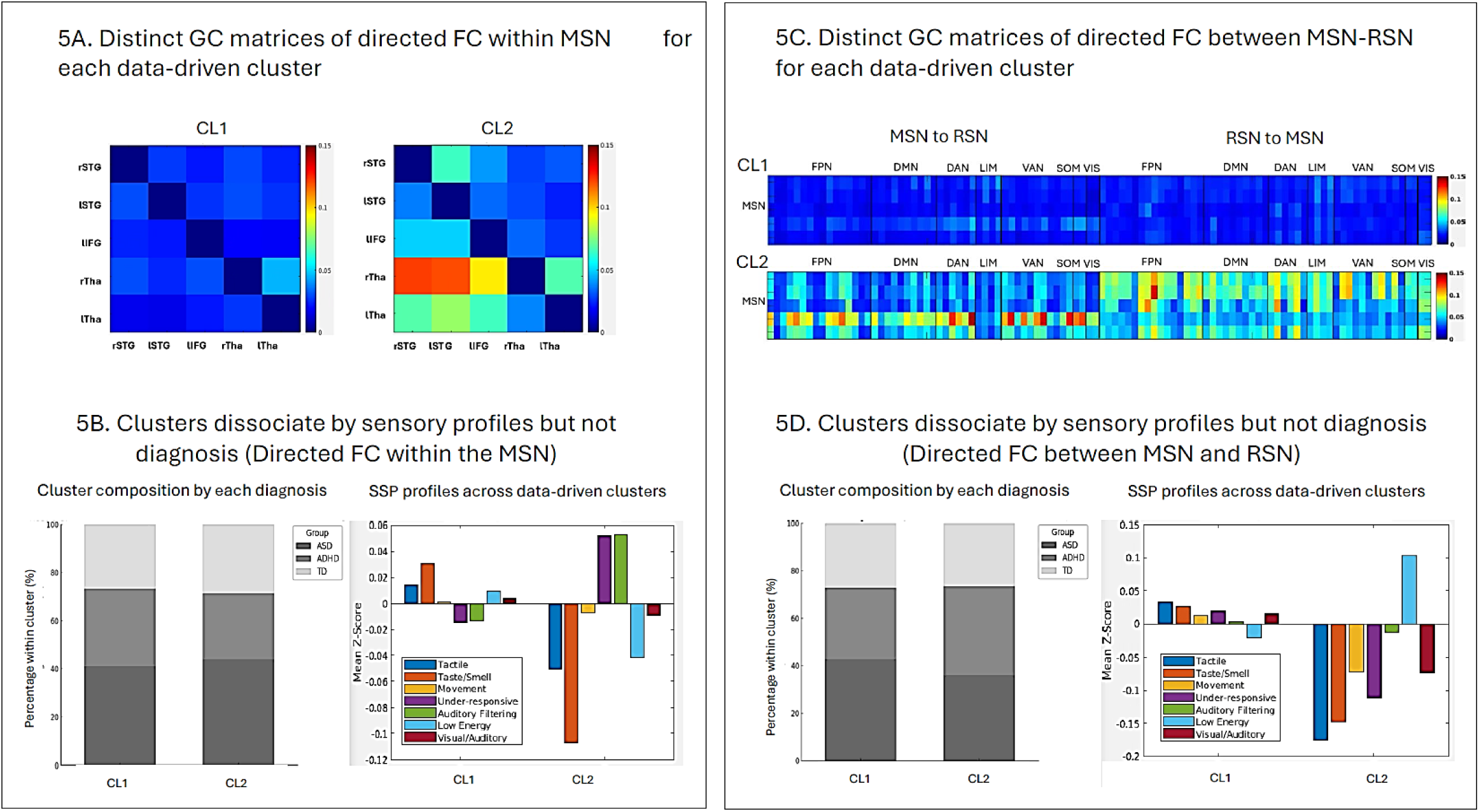
Data-driven clusters based on directed FC within the MSN and between the MSN and RSNs. **5A**. Two data-driven clusters exhibited distinct patterns of directed FC within the MSN, **5B**. When normalized by cluster size (%), both clusters included participants from each diagnostic group, but demonstrated distinct sensory profile features: Cluster 1 (mild sensory issues for under-responsive and auditory filtering), Cluster 2 (severe sensory issues particularly taste and smell), **5C**. Two data-driven clusters also showed distinct directed FC patterns between the MSN and RSNs, **5D**. When normalized by cluster size (%), both clusters again included all diagnostic groups, but demonstrated distinct sensory profiles: Cluster 1 (sensory adaptive with mild issue with low energy), Cluster 2 (generalized sensory issues except low energy) GC, Granger Causality; rSTG, Right Superior Temporal Gyrus; lSTG, Left Superior Temporal Gyrus; lIFG, Left Inferior Frontal Gyrus; rTha, Right Thalamus; lTha, Left Thalamus; FPN, Fronto-parietal network; DMN, Default-mode network; DAN, Dorsal attention network; LIM, Limbic network; VAN, Ventral attention network; SOM, Somatomotor network; VIS, Visual network

Similarly, the between MSN-RSN FC analysis also identified two optimal clusters (Supplementary Figure 4B), with distinct FC patterns (**Figure 5**C). The FC strength across MSN-RSN connections increased in Cluster2 (N=64) compared to Cluster1 (N=353), notably greater directed FC from right Thalamus to RSNs. The clusters did not significantly differ by age (t=1.46, p=0.15), sex (χ²=1.59, p=0.45) and scanner (χ²=0.12, p=0.94). Within each diagnostic group, the two clusters were proportionally distributed (Cluster1: Cluster2): TD=85%, 15%; ASD=86.8%, 12.2%; ADHD=81.5%, 18.5% (χ²=1.59, p=0.45). Notably, sensory profiles were more distinct across clusters than diagnostic distributions (**Figure 5**D).

## Discussion

The present study investigated the functional organization of the MSN at resting-state by examining both within-MSN and between MSN-RSN FC and their relations to both diagnosis and clinical sensory characteristics. Overall, thalamus and temporal regions exhibited the strongest functional connectivity both within the MSN and between MSN and other RSNs. Importantly, we found fewer diagnostic differences in FC both within the MSN and between MSN and other RSNs. Instead, differences in FC were strongly associated with clinical sensory issues themselves, regardless of diagnoses. These results suggest that the underlying neurobiology in NDDs, despite diagnostic differences in parent-reported sensory issues, are transdiagnostic in nature as opposed to specific to ASD or ADHD.

In terms of the MSN broadly, the bilateral STG, MTG, and the right insula exhibited strong connectivity both within the MSN and with somatomotor and visual networks in the RSNs. The thalamus demonstrated robust directional influence not only toward other MSN subregions, but also across nearly all RSN subregions, beyond sensory networks, extending into higher-order association networks. This finding aligns with recent reviews highlighting the thalamic pulvinar as a critical hub for integrating and regulating sensory information across modalities.^53^ The pulvinar facilitates hierarchical sensory processing through its extensive reciprocal connections with lower-order sensory regions and higher-order associative cortical areas, enabling effective integration and coordination of multisensory information.^54–57^

The overall functional organization of the MSN was largely shared between individuals with ASD and ADHD, with only a single connection showing differences within the MSN. However, the ADHD group showed more consistently elevated FC compared to TD group, while the ASD group exhibited more heterogeneous FC patterns. Diagnostic differences of ASD and ADHD relative to TD in MSN functional organization were more pronounced in FC between the MSN and RSNs than within the MSN itself, suggesting atypical MSN organization in neurodevelopmental conditions at a large-scale network level in associations with other RSNs.

Both ASD and ADHD groups exhibited overall increased connectivity strength between the MSN and RSNs, particularly with the DMN, compared to the TD group. Notably, this atypical MSN-DMN organization was further characterized by a reduction in directional influences from MSN to DMN and increased directional influences from DMN to MSN in both clinical groups. These findings suggest transdiagnostic alterations in how multisensory information is integrated with higher-order cognitive processes. While the DMN is traditionally associated with self-referential thinking and mind-wandering, and its atypical functional organization has been reported across various psychiatric and neurodevelopmental conditions,^58,59^ it also plays a critical role in dynamically interacting with multisensory regions to integrate and modulate sensory information, particularly when conflicts or ambiguities arise.^60,61^ The altered MSN-DMN organization observed in both clinical groups may reflect a shift in functional hierarchy, where MSN’s influence over DMN is diminished, while DMN exerts a stronger influence on sensory integration. This shift may reflect altered sensory integration processes that could underlie difficulties in adapting to dynamic sensory environments. Moreover, this hierarchical reorganization aligns with previous findings of altered network integration and segregation in ASD and ADHD, where disrupted organization between task-free and task-positive networks can lead to difficulties in goal-directed behaviour and multisensory processing.^62,63^ The shared pattern across ASD and ADHD suggests a potential common developmental mechanism, such as inefficient modulation of sensory input by higher-order networks.^62^

Compared to the ASD group, which exhibited more complex patterns of connectivity differences, the ADHD group distinctively showed increased connectivity strength between the MSN and multiple RSNs involved in cognitive control, attention, and sensorimotor processing while controlling for age and sex. In the directional influences, the rSTG consistently emerged as a key node in the altered organization of the MSN. While both clinical groups commonly exhibited decreased directional influences from MSN to RSNs and increased influences from RSNs to MSN, ADHD showed a distinct, opposite pattern, characterized by increased directional influence from rSTG to FPN and decreased influence from FPN/DAN to rSTG. These findings may reflect a condition-specific compensatory mechanism in ADHD involving STG, characterized by enhanced bottom-up connectivity from multisensory regions to higher-order networks, alongside reduced top-down influences from frontoparietal and dorsal attentional network to the rSTG. This imbalance could contribute to difficulties in attentional control and sensory regulation commonly observed in ADHD,^64,65^ while the broader and more variable MSN-RSN alterations observed in ASD support high heterogeneity within this group.^66,67^

Beyond the diagnostic differences, the findings supported that MSN organization strongly aligns with sensory processing difficulties across all participants. Increased connectivity strength between the bilateral thalami and other MSN sub-regions within the MSN, as well as between the MSN and other RSNs involving nearly all MSN sub-regions, was strongly associated with poorer daily sensory processing abilities. Increased connectivity strength between the MSN and RSNs may reflect reduced network segregation, leading to inefficiencies in filtering, prioritizing, and responding appropriately to sensory input. This global disorganization could manifest as heightened sensory sensitivity, sensory seeking, or under-responsiveness. This aligns with models suggesting that sensory over-connectivity can lead to heightened sensory reactivity and reduced adaptability to dynamic environments.^68,69^

In contrast, the associations between directed connectivity and daily sensory processing abilities were more complex and less robust than those observed with connectivity strength. Since the MSN primarily processes incoming sensory inputs while RSNs modulate attention, cognition, and self-regulation, daily sensory processing abilities may be more influenced by the overall co-activation and balance of these networks rather than by a strict top-down or bottom-up directional hierarchy. Notably, the majority of the findings in the directed connectivity involved the thalamus. Given that the thalamus played a role as a sensory relay hub in shaping MSN directional organization across all participants, independent of diagnosis, altered thalamic function may contribute to inefficient sensory gating and filtering, amplifying sensory difficulties across multiple domains.^70–72^

Most strikingly, our data-driven approaches revealed that heightened FC indices in both connectivity strength and directional influence were associated with greater sensory processing difficulties, regardless of diagnostic labels. Notably, increased connectivity strength was broadly linked to generalized sensory processing difficulties, while increased directional influence appeared to be associated with challenges in more specific sensory domains or features.

Specifically, the group characterized by heightened directional influence from the thalamus within the MSN showed pronounced difficulties related to taste/smell and tactile sensitivities. In contrast, heightened directional influence from the thalamus between the MSN and other RSNs was associated with generalized sensory difficulties. These findings suggest that the flow of information within and between networks may be more relevant for certain sensory modalities. Moreover, the association between generalized sensory difficulties and heightened directional influence from the thalamus between the MSN and RSNs suggests that sensory regulation may rely on the interaction between sensory and large-scale networks. Excessive cross-network influence could contribute to sensory overload or inefficient sensory filtering.

In sum, our data suggests that the functional organization of the MSN reflects a distributed and integrative structure, characterized by strong connections and bidirectional influence between lower-order and higher-order brain regions and networks. The maturation of the MSN is reflected in decreasing overall connectivity strength and directional influence indices, supporting greater efficiency. However, the ASD and ADHD groups demonstrated increased patterns across multiple aspects, highlighting their divergent and atypical patterns compared to their typically developing peers. The ADHD group showed a more distinctive pattern involving the STG, particularly in its connections with fronto-parietal control, attention, and sensorimotor regions. In contrast, the ASD group exhibited broader heterogeneous patterns, involving not only higher-order regions and networks, such as the STG, IFG, and DMN, but also lower-order structures and networks, including the thalamus and limbic system. While we identified some diagnosis-specific differences in the functional organization of the MSN, sensory processing abilities - independent of diagnostic categories - showed stronger and more pervasive associations with all aspects of MSN functional organization. Data-driven clustering further confirmed that varying levels of connectivity strength and directional influence align more closely with sensory processing abilities than with diagnostic labels. These findings suggest that the functional organization of the MSN is intricately tied to daily sensory processing abilities, emphasizing its fundamental role of integrating and regulating sensory inputs not only within the MSN but also with dynamic interactions with other large-scale networks.

## Acknowledgements

This work was directly funded by Ontario Brain Institute, and a CIHR Project Grant to RAS, BS, EA, and AK (487850). RAS is funded through a Dorothy Killam Fellowship, an NSERC Discovery Grant (RGPIN-2024-06233), two SSHRC Insight Grants (435-2017-0936 & 435-2024-1375), the University of Western Ontario Faculty Development Research Fund, and a Canadian Foundation for Innovation John R. Evans Leaders Fund (37497), and through a grant from the Canada First Research Excellence Fund (BrainsCAN). RS would also like to thank Eddie V., Stone S., Jeff A., Dave K., and Mike M. for their inspirational work from 1991 tackling social concerns including childhood trauma and mental health.

## CRediT Statement

EJC – Conceptualization, methodology, formal analysis, data curation, writing – original draft, writing – review & editing. KL–Methodology, writing – review & editing. MV–Methodology, writing – review & editing. EA – Data collection, investigation, writing – review & editing. PA – Data collection, investigation, writing – review & editing. MA– Data collection, investigation, writing – review & editing. JC– Data collection, investigation, writing – review & editing. SG– Data collection, investigation, writing – review & editing. JJ– Data collection, investigation, writing – review & editing. EK– Data collection, investigation, writing – review & editing. AK– Data collection, investigation, writing – review & editing. JPL– Data collection, investigation, writing – review & editing. RS– Data collection, investigation, writing – review & editing. BS– Conceptualization, methodology, investigation, writing – review & editing, funding acquisition. MJT– Data collection, investigation, writing – review & editing. RS – Conceptualization, methodology, software, formal analysis, investigation, resources, data curation, writing – review and editing, supervision, project administration, funding acquisition.

## Supplementary information

### Data preprocessing

All imaging data were preprocessed using *fMRIPrep*^1^, a pipeline built upon the Nipype^2^, incorporating both anatomical and functional workflows. For anatomical preprocessing, T1-weighted structural images underwent bias field correction to address intensity non-uniformity^3^, followed by skull stripping using the OASIS template and Advanced Normalization Tools (ANTs) to generate brain masks. Surface reconstruction was performed with FreeSurfer^4^, and the resulting cortical surfaces were used to refine the brain masks through a customized version of the Mindboggle^5^. Nonlinear registration using ANTs^6^ then aligned each participant’s skull-stripped image to a pediatric brain template^7,8^. Subsequently, FSL tools were used to segment the normalized anatomical images into cerebrospinal fluid (CSF), white matter (WM), and gray matter compartments^9^. For functional preprocessing, data were corrected for slice timing and motion using AFNI^10^ and FSL^11^, respectively. To correct for susceptibility distortions in the absence of fieldmaps, each participant’s functional image was co-registered to an inverted T1-weighted structural image using a fieldmap-template-constrained ANTs registration^12–14^. Coregistration between functional and structural images was refined using boundary-based registration with six degrees of freedom via FreeSurfer^15^. The transformations from motion correction, functional-to-anatomical alignment, and anatomical-to-template normalization were combined and applied in one step using ANTs with Lanczos interpolation. Head motion metrics were calculated using Nipype’s implementation, including framewise displacement (FD)^16^ and DVARS (the standardized derivative of root mean square variance over voxels^17^. Participants exceeding the exclusion threshold of more than one-third of frames with FD > 0.5 mm or DVARS > 1.5 were excluded from subsequent analyses. Following fMRIPrep preprocessing, additional denoising was performed. Six motion parameters along with signal components from WM and CSF (including their temporal derivatives and quadratic terms) were regressed from the data. High-pass temporal filtering with a cutoff of 0.008 Hz was applied concurrently using AFNI^10,17^.

### Short Sensory Profile (SSP)

The Short Sensory Profile (SSP) is a widely used and well-validated 38-item parent-report measure designed to assess behaviours linked to atypical responses to sensory input in children aged 3 to 10 years^18–20^. This questionnaire evaluates sensory processing across seven domains: tactile sensitivity (7 items), taste/smell sensitivity (4 items), movement sensitivity (3 items), under-responsiveness/seeking sensation (7 items), auditory filtering (6 items), low energy/weakness (6 items), and visual/auditory sensitivity (5 items). Parents rate each item on a 5-point Likert scale, where 1 indicates “always” (100% of the time), 2 “frequently” (75%), 3 “occasionally” (50%), 4 “seldom” (25%), and 5 “never” (0%), reflecting how often the child exhibits each behaviour. Lower scores in the total SSP score indicate greater sensory processing difficulties. The SSP has demonstrated robust psychometric properties, including strong internal consistency in autistic populations (Cronbach’s alpha = 0.89)^18^ and reliable discriminative validity in differentiating children with and without sensory processing challenges. While originally developed for typically developing children, the SSP’s validity has been established through its consistent associations with clinical diagnoses and observational measures of sensory function in both community and clinical samples^21,22^.

---

**Supplementary Table1.**
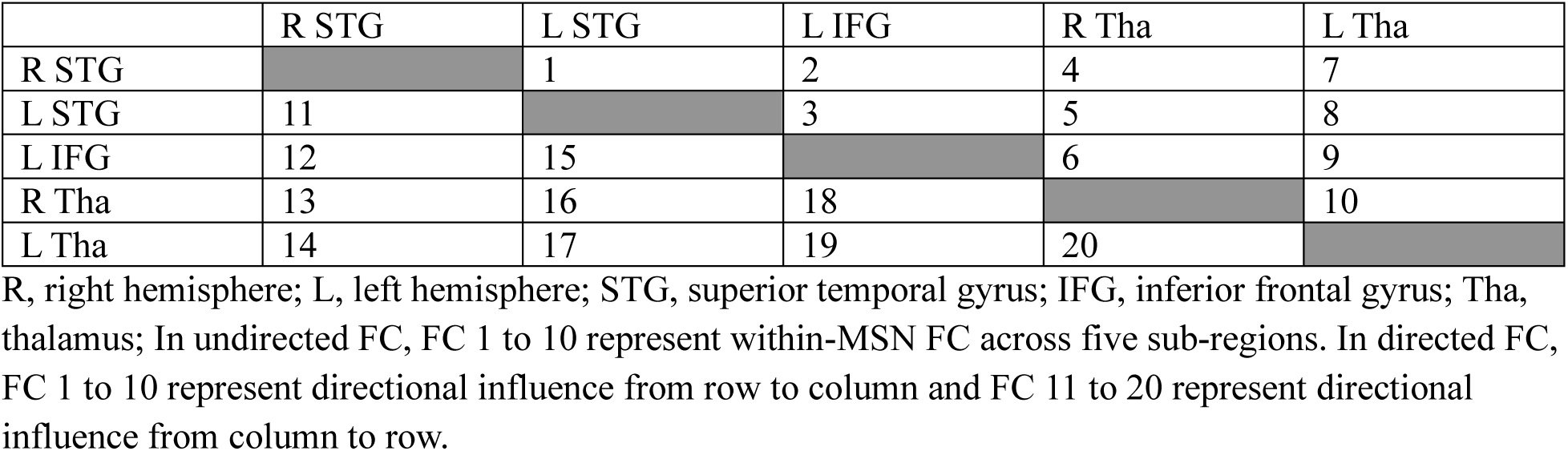
Definition of Multisensory Network (MSN) connectivity based on ROI pairings

**Supplementary Table2.**
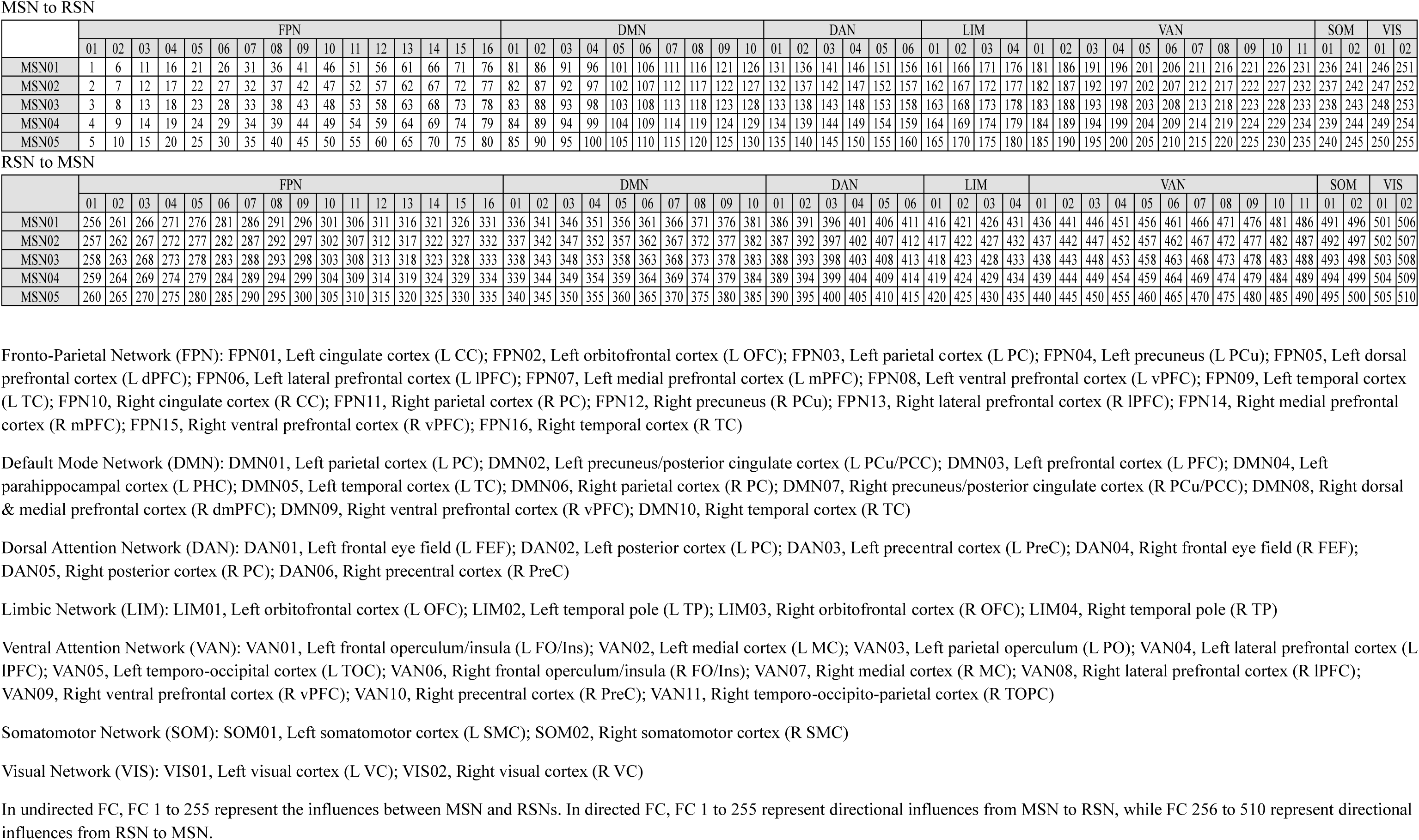
Definition of connectivity between Multisensory Network (MSN) and Resting-state Networks (RSNs) based on ROI pairings

**Supplementary Table 3.**
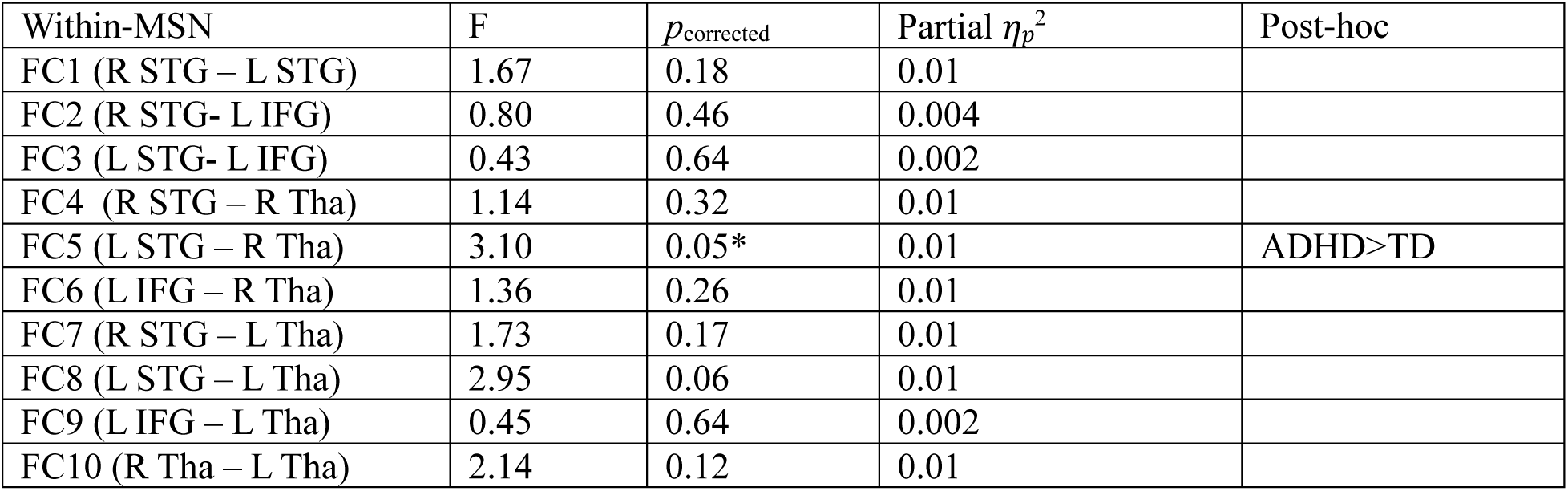
Diagnostic differences in the undirected FC within-MSN while controlling for age and sex

**Supplementary Table 4.**
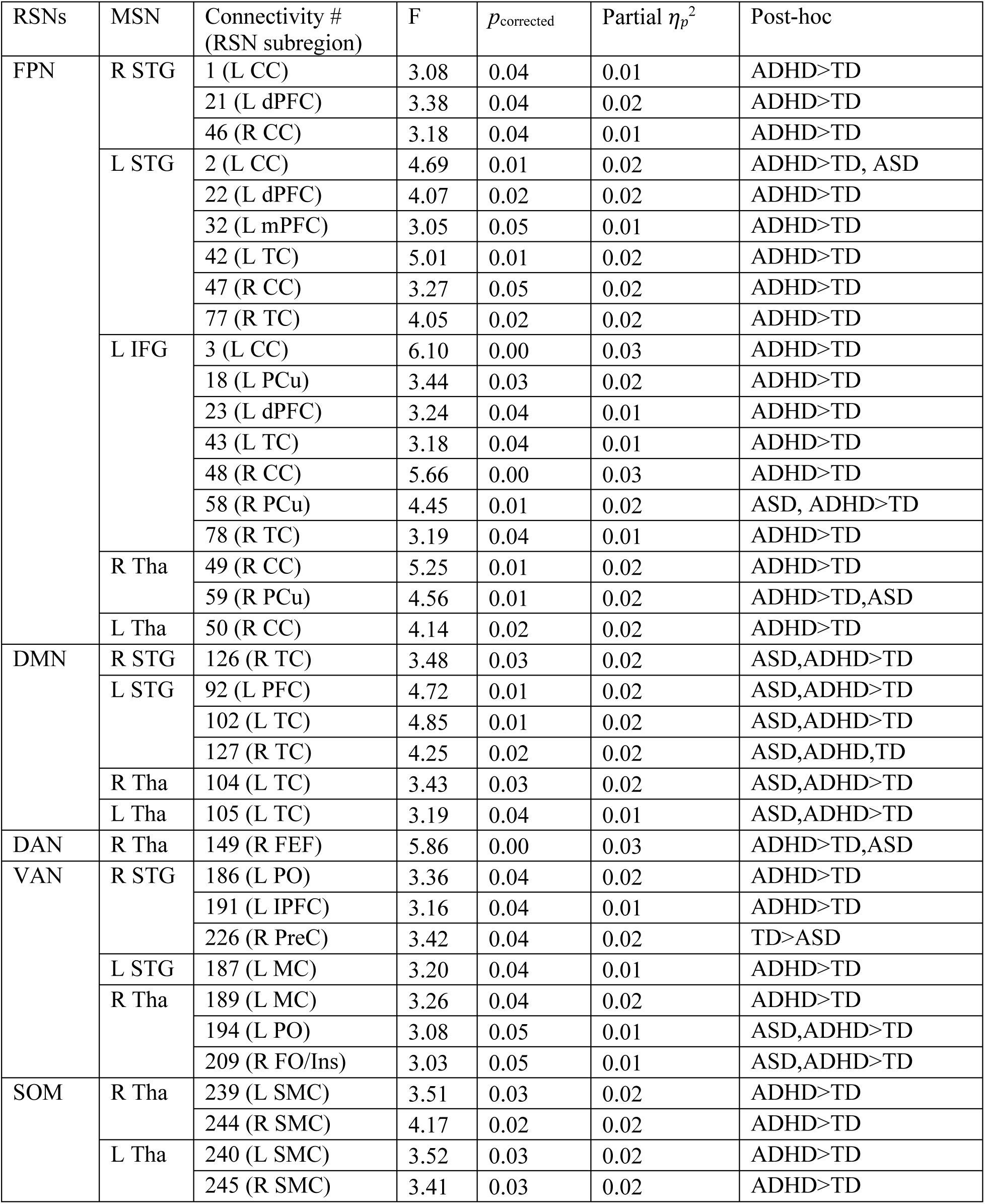
Diagnostic differences in the undirected FC between MSN-RSN while controlling for age and sex (only FCs identified as significant at *p*_corrected_<.05 are presented)

**Supplementary Table 5.**
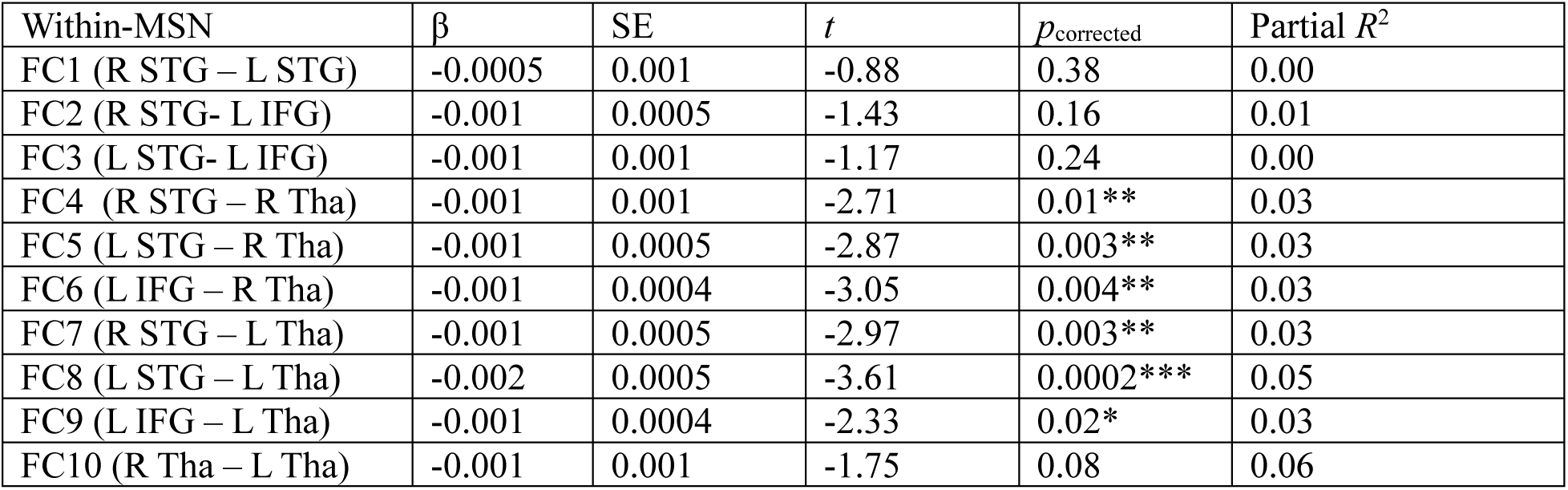
Associations between sensory processing abilities and the undirected FC within-MSN while controlling for age and sex

**Supplementary Table 6.**
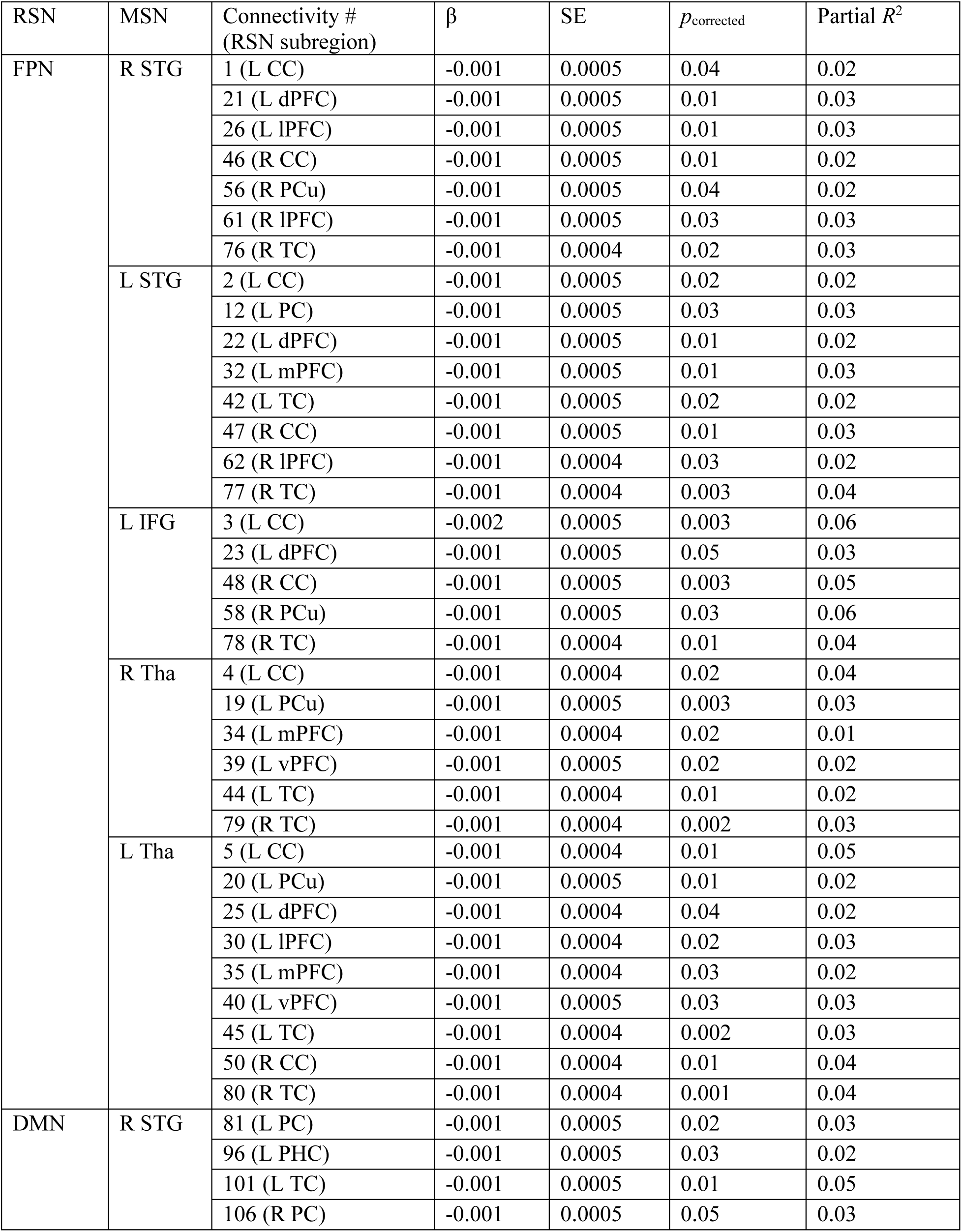
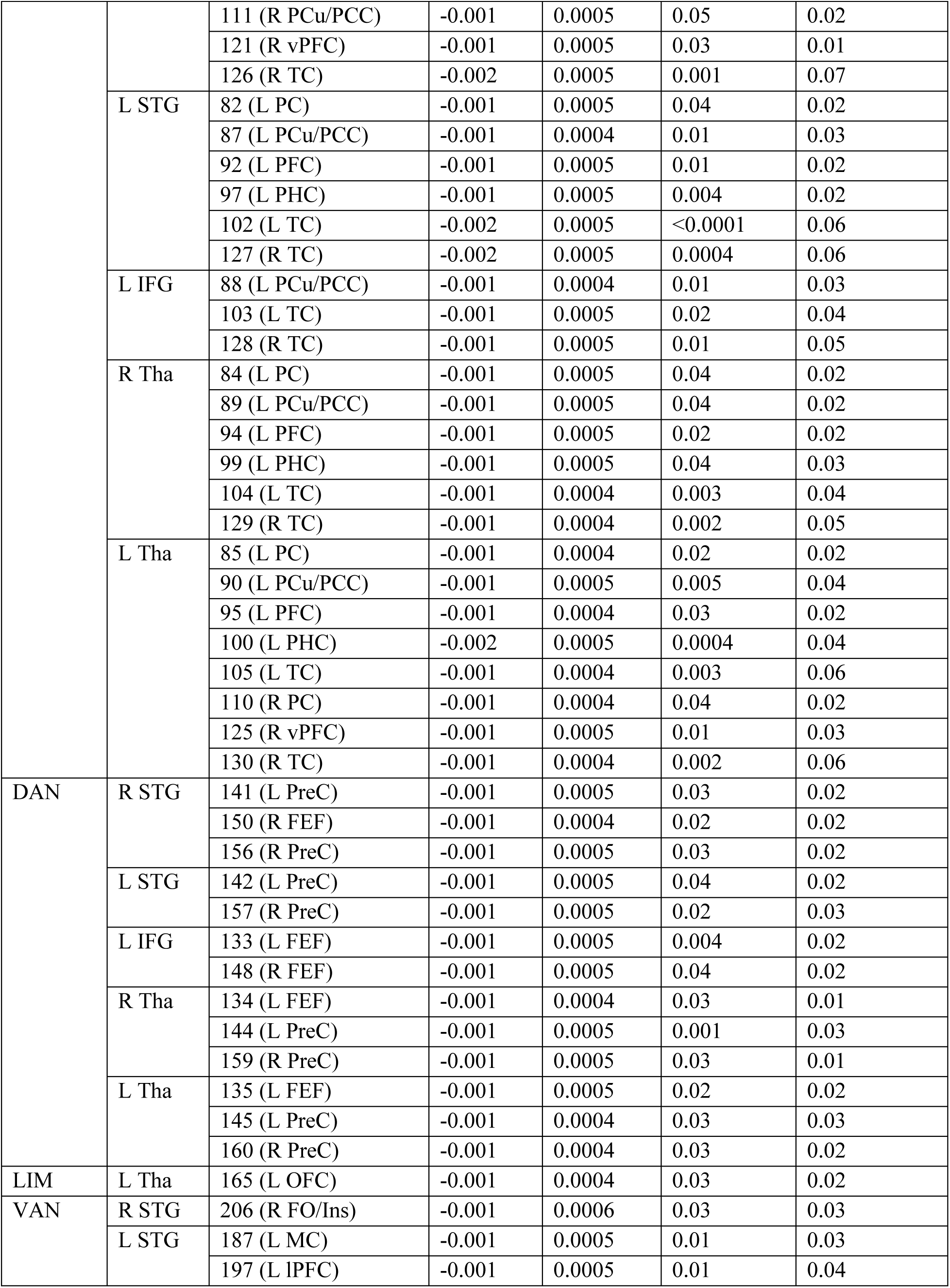
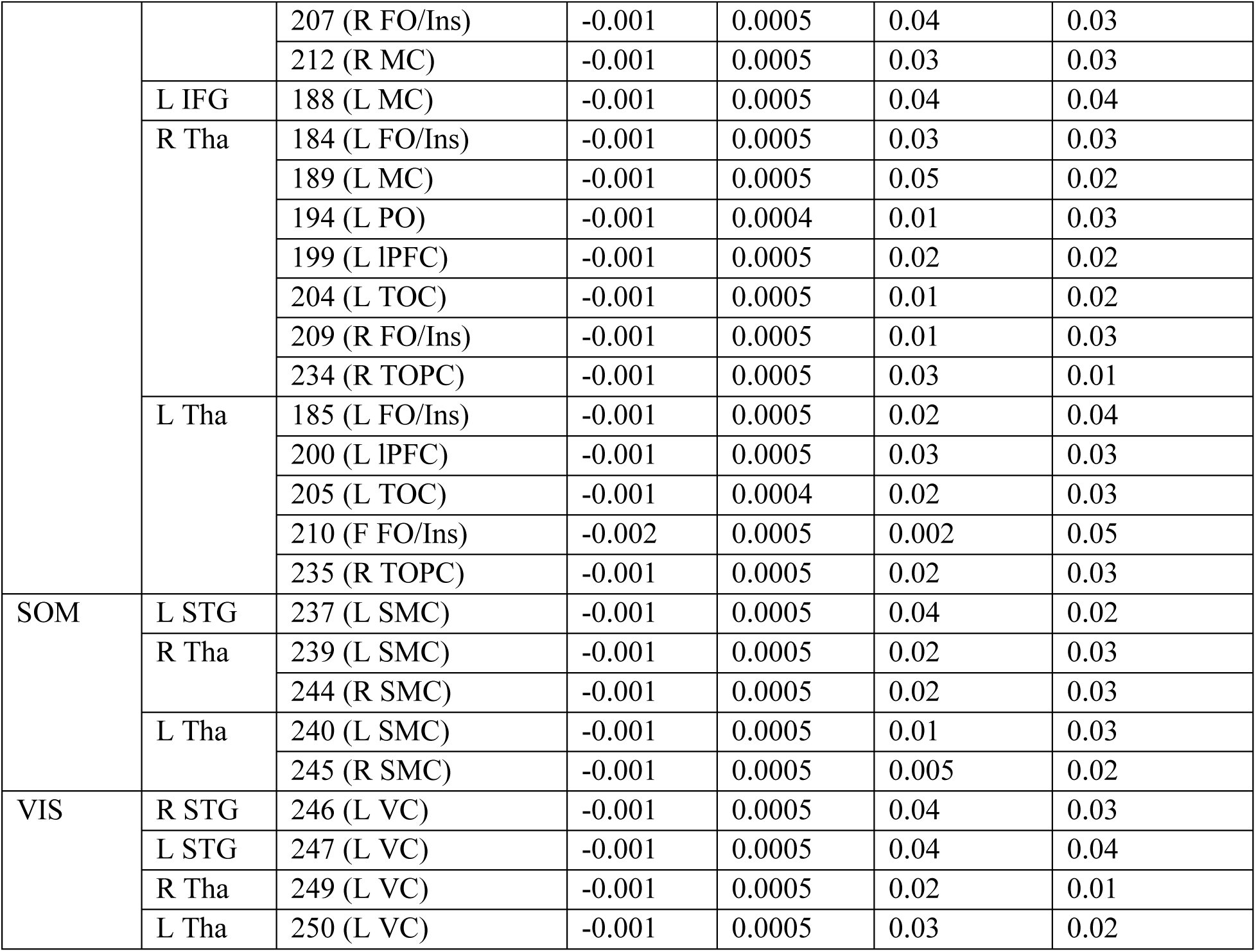
Associations between sensory processing abilities and the undirected FC within-MSN while controlling for age and sex (only FCs identified as significant at *p*_corrected_ <.05 are presented)

**Supplementary Table 7.**
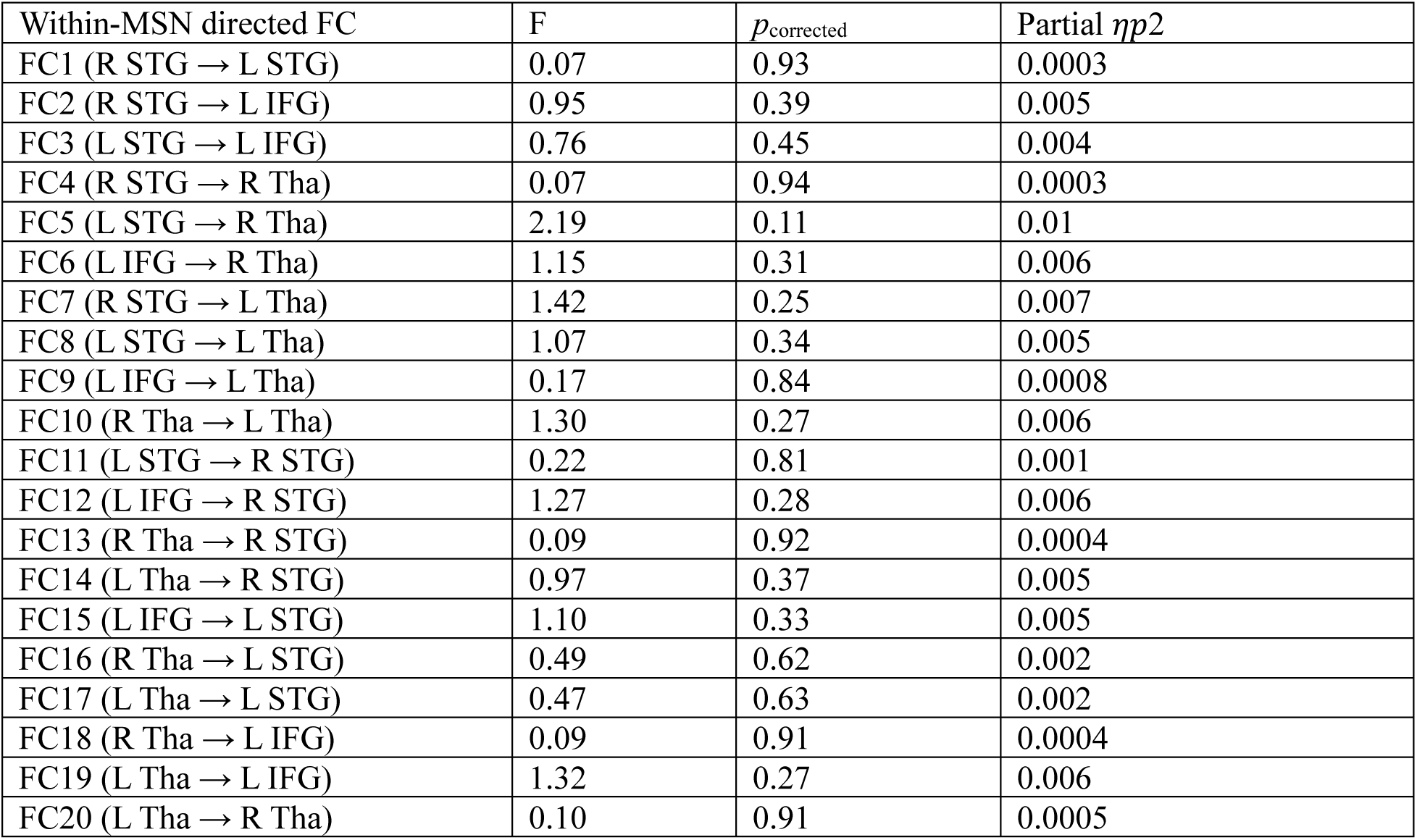
Diagnostic differences in the directed FC within-MSN while controlling for age and sex

| Within-MSN directed FC | F | $p_{\text{corrected}}$ | Partial $\eta p^2$ |
| --- | --- | --- | --- |
| FC1 (R STG → L STG) | 0.07 | 0.93 | 0.0003 |
| FC2 (R STG → L IFG) | 0.95 | 0.39 | 0.005 |
| FC3 (L STG → L IFG) | 0.76 | 0.45 | 0.004 |
| FC4 (R STG → R Tha) | 0.07 | 0.94 | 0.0003 |
| FC5 (L STG → R Tha) | 2.19 | 0.11 | 0.01 |
| FC6 (L IFG → R Tha) | 1.15 | 0.31 | 0.006 |
| FC7 (R STG → L Tha) | 1.42 | 0.25 | 0.007 |
| FC8 (L STG → L Tha) | 1.07 | 0.34 | 0.005 |
| FC9 (L IFG → L Tha) | 0.17 | 0.84 | 0.0008 |
| FC10 (R Tha → L Tha) | 1.30 | 0.27 | 0.006 |
| FC11 (L STG → R STG) | 0.22 | 0.81 | 0.001 |
| FC12 (L IFG → R STG) | 1.27 | 0.28 | 0.006 |
| FC13 (R Tha → R STG) | 0.09 | 0.92 | 0.0004 |
| FC14 (L Tha → R STG) | 0.97 | 0.37 | 0.005 |
| FC15 (L IFG → L STG) | 1.10 | 0.33 | 0.005 |
| FC16 (R Tha → L STG) | 0.49 | 0.62 | 0.002 |
| FC17 (L Tha → L STG) | 0.47 | 0.63 | 0.002 |
| FC18 (R Tha → L IFG) | 0.09 | 0.91 | 0.0004 |
| FC19 (L Tha → L IFG) | 1.32 | 0.27 | 0.006 |
| FC20 (L Tha → R Tha) | 0.10 | 0.91 | 0.0005 |

**Supplementary Table 8.**
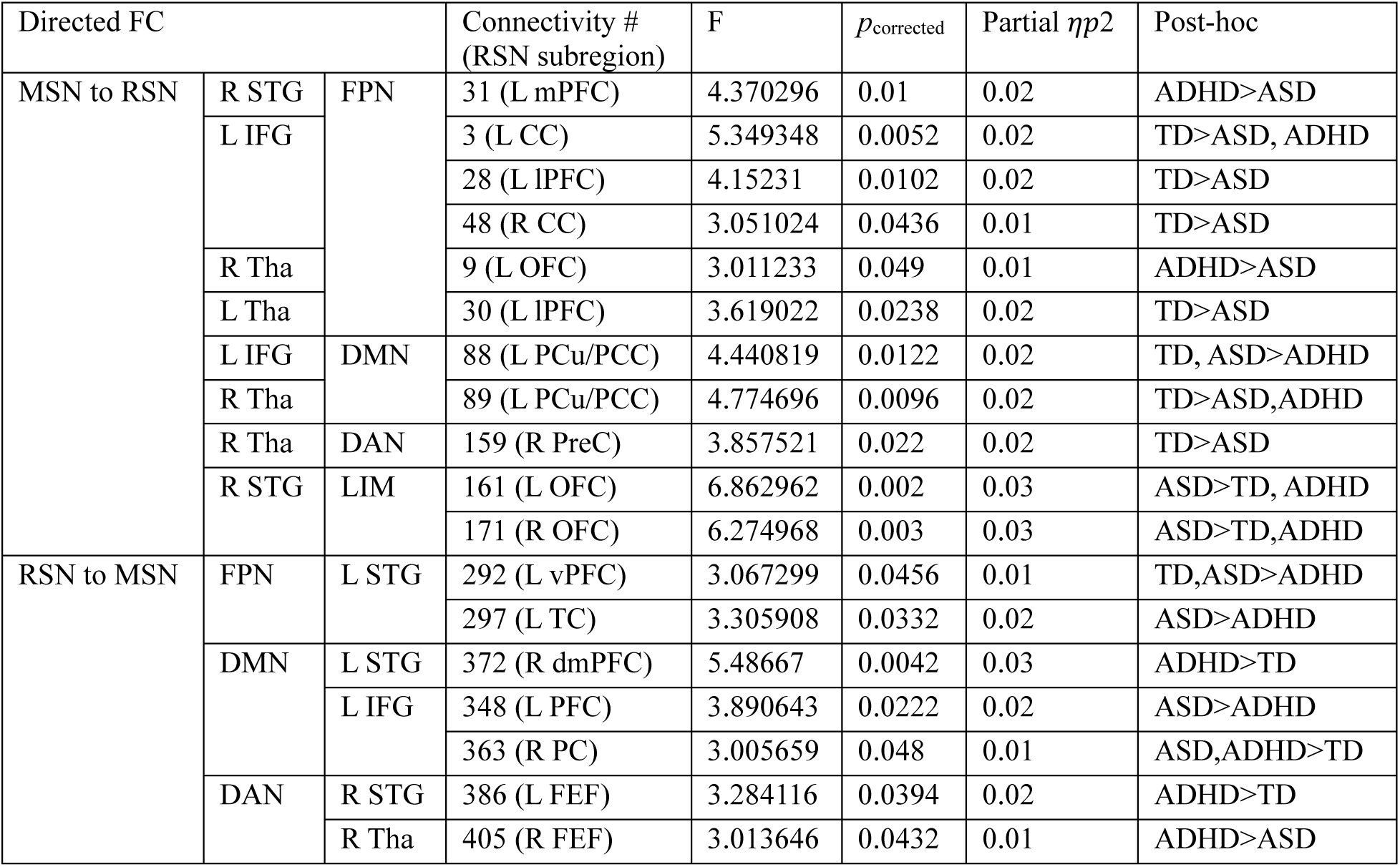
Diagnostic differences in the directed FC between MSN-RSN while controlling for age and sex (only FCs identified as significant at *p*_corrected_ <.05 are presented)

**Supplementary Table 9.**
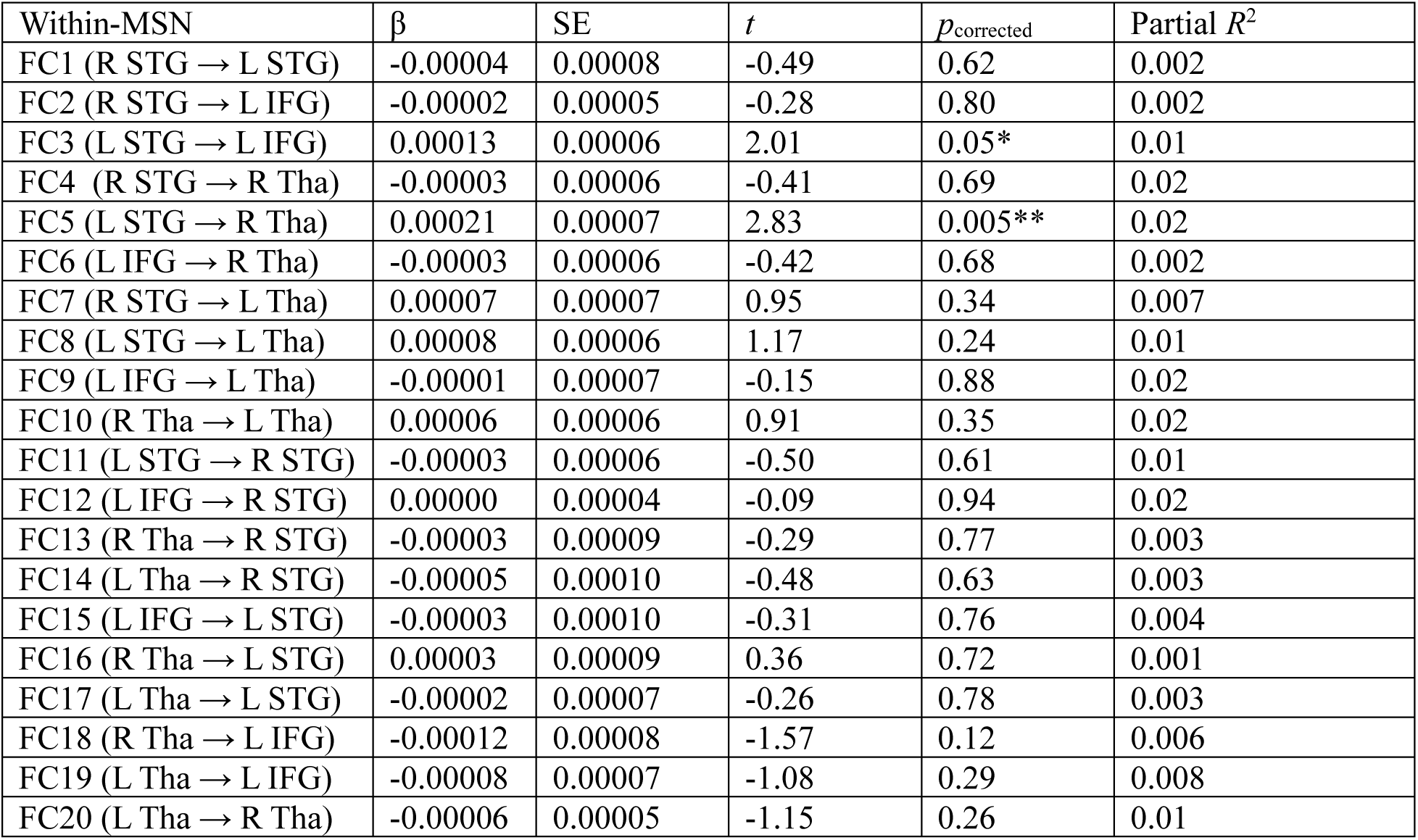
Associations between sensory processing abilities and the directed FC within-MSN while controlling for age and sex

| Within-MSN | $\beta$ | SE | $t$ | $p_{\text{corrected}}$ | Partial $R^2$ |
| --- | --- | --- | --- | --- | --- |
| FC1 (R STG → L STG) | -0.00004 | 0.00008 | -0.49 | 0.62 | 0.002 |
| FC2 (R STG → L IFG) | -0.00002 | 0.00005 | -0.28 | 0.80 | 0.002 |
| FC3 (L STG → L IFG) | 0.00013 | 0.00006 | 2.01 | 0.05* | 0.01 |
| FC4 (R STG → R Tha) | -0.00003 | 0.00006 | -0.41 | 0.69 | 0.02 |
| FC5 (L STG → R Tha) | 0.00021 | 0.00007 | 2.83 | 0.005** | 0.02 |
| FC6 (L IFG → R Tha) | -0.00003 | 0.00006 | -0.42 | 0.68 | 0.002 |
| FC7 (R STG → L Tha) | 0.00007 | 0.00007 | 0.95 | 0.34 | 0.007 |
| FC8 (L STG → L Tha) | 0.00008 | 0.00006 | 1.17 | 0.24 | 0.01 |
| FC9 (L IFG → L Tha) | -0.00001 | 0.00007 | -0.15 | 0.88 | 0.02 |
| FC10 (R Tha → L Tha) | 0.00006 | 0.00006 | 0.91 | 0.35 | 0.02 |
| FC11 (L STG → R STG) | -0.00003 | 0.00006 | -0.50 | 0.61 | 0.01 |
| FC12 (L IFG → R STG) | 0.00000 | 0.00004 | -0.09 | 0.94 | 0.02 |
| FC13 (R Tha → R STG) | -0.00003 | 0.00009 | -0.29 | 0.77 | 0.003 |
| FC14 (L Tha → R STG) | -0.00005 | 0.00010 | -0.48 | 0.63 | 0.003 |
| FC15 (L IFG → L STG) | -0.00003 | 0.00010 | -0.31 | 0.76 | 0.004 |
| FC16 (R Tha → L STG) | 0.00003 | 0.00009 | 0.36 | 0.72 | 0.001 |
| FC17 (L Tha → L STG) | -0.00002 | 0.00007 | -0.26 | 0.78 | 0.003 |
| FC18 (R Tha → L IFG) | -0.00012 | 0.00008 | -1.57 | 0.12 | 0.006 |
| FC19 (L Tha → L IFG) | -0.00008 | 0.00007 | -1.08 | 0.29 | 0.008 |
| FC20 (L Tha → R Tha) | -0.00006 | 0.00005 | -1.15 | 0.26 | 0.01 |

**Supplementary Table 10.**
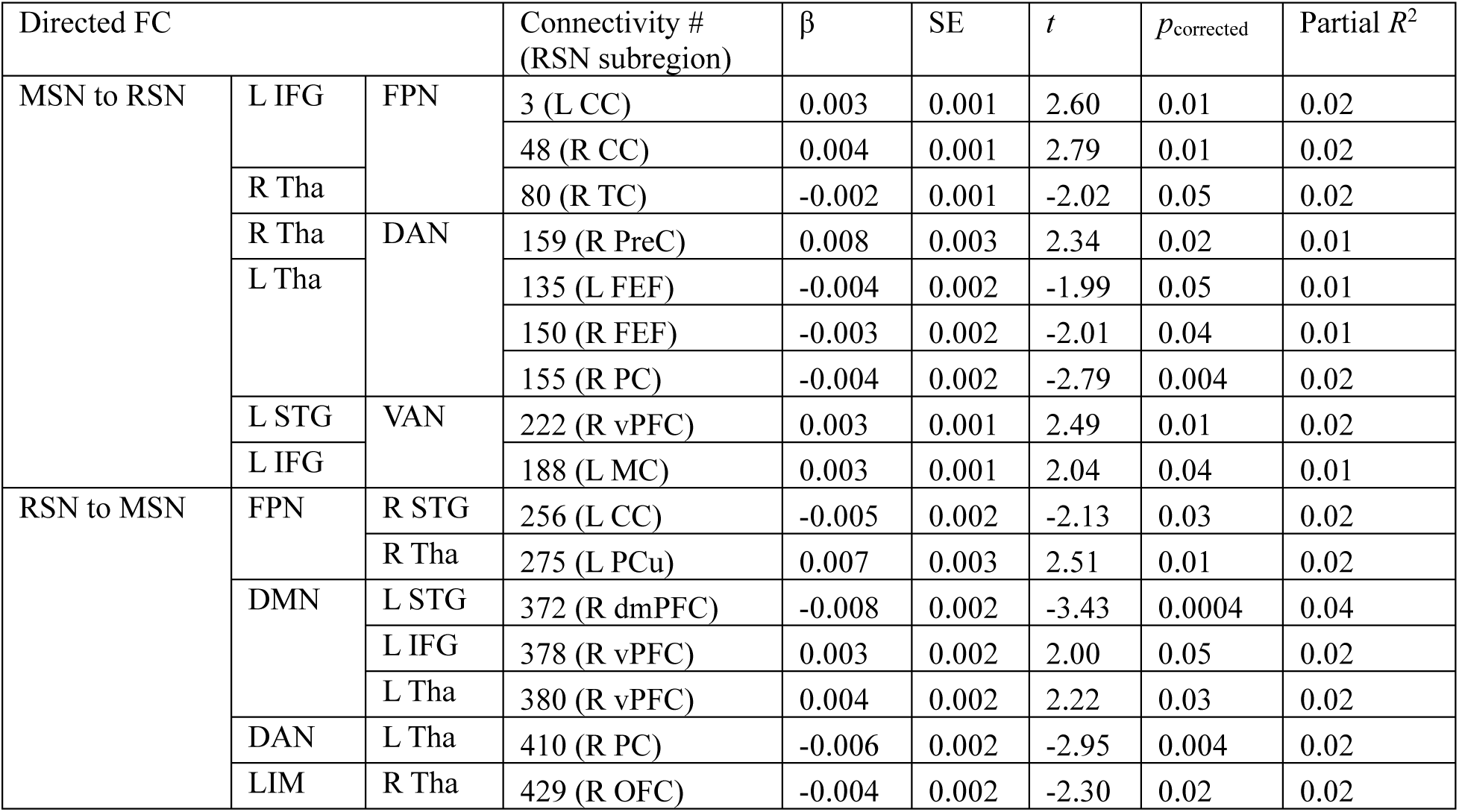
Associations between sensory processing abilities and the directed FC between MSN-RSN while controlling for age and sex (only FCs identified as significant at *p*_corrected_<.05 are presented)

**Supplementary Figure 1.**
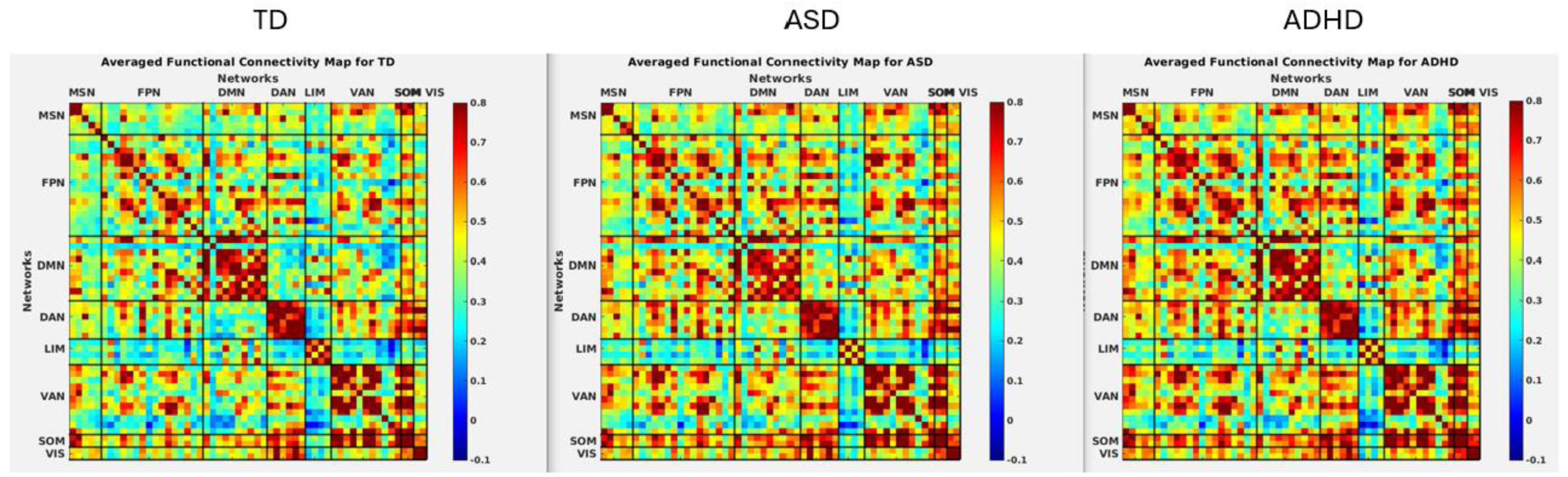
Undirected functional connectivity maps within and between MSN and RSNs in three groups.

**Supplementary Figure 2.**
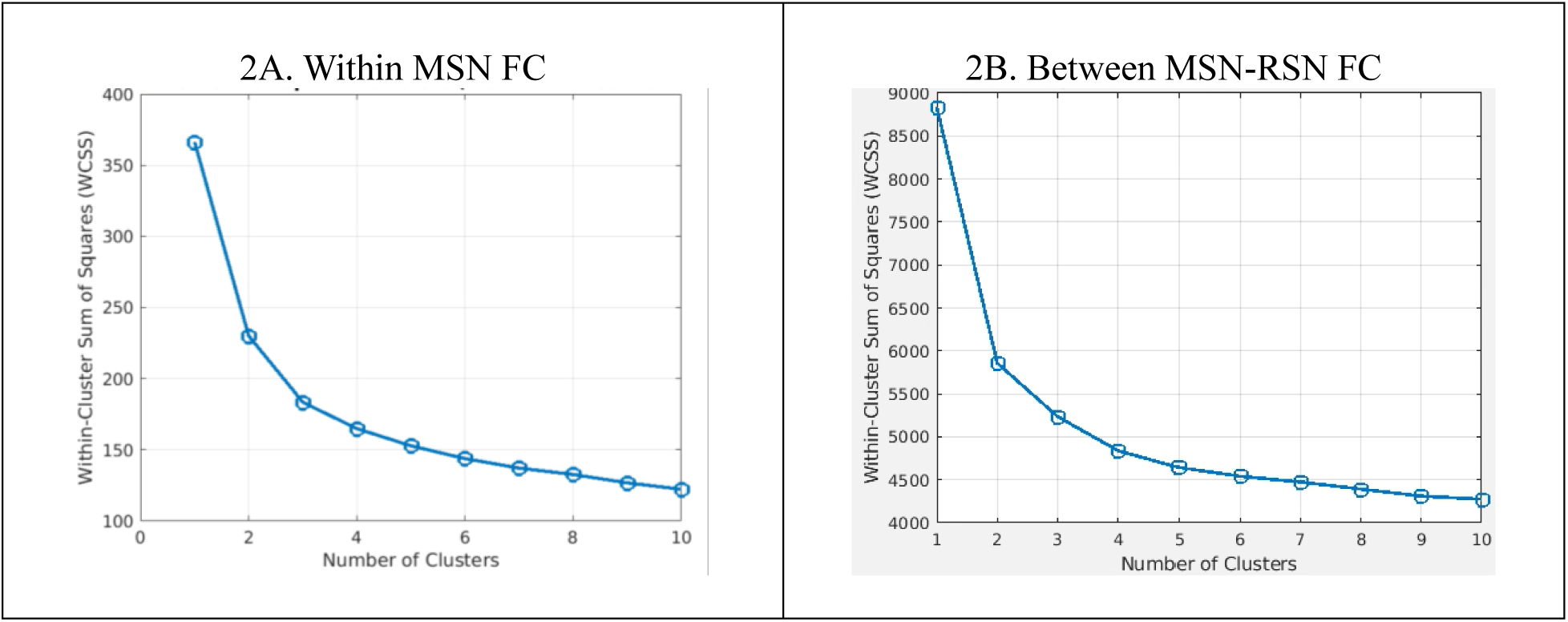
The optimal numbers of clusters in the undirected FC within-MSN (2A) and between MSN-RSNs (2B) from elbow method

**Supplementary Figure 3.**
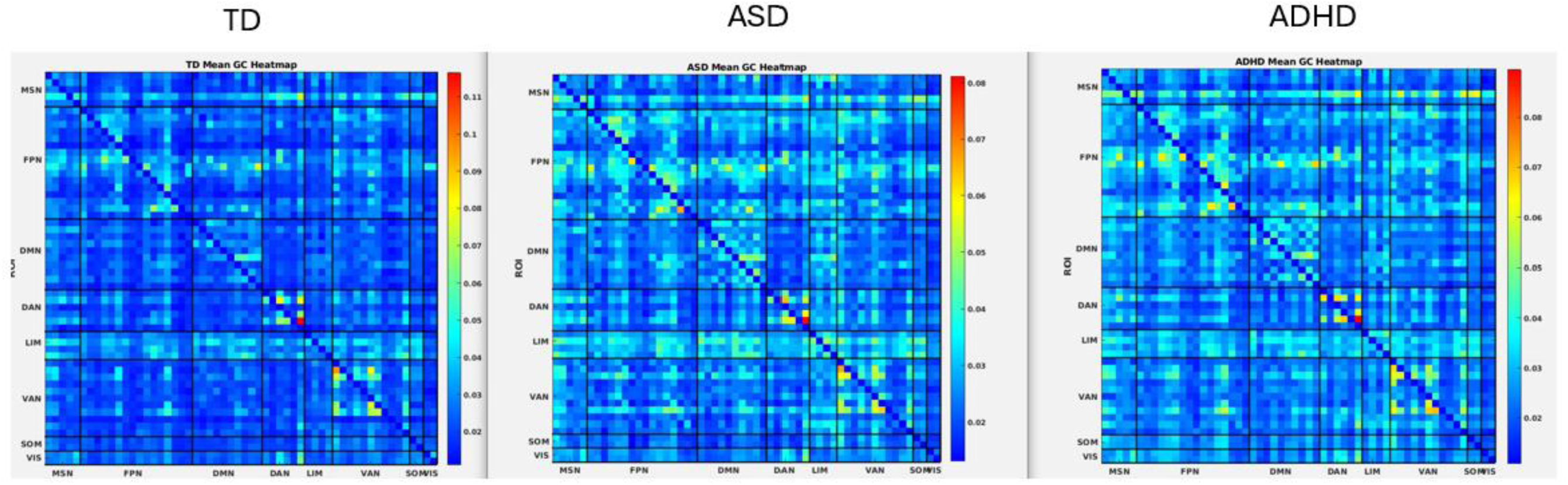
Directed functional connectivity maps within MSN and between MSN and RSNs in three groups.

**Supplementary Figure 4.**
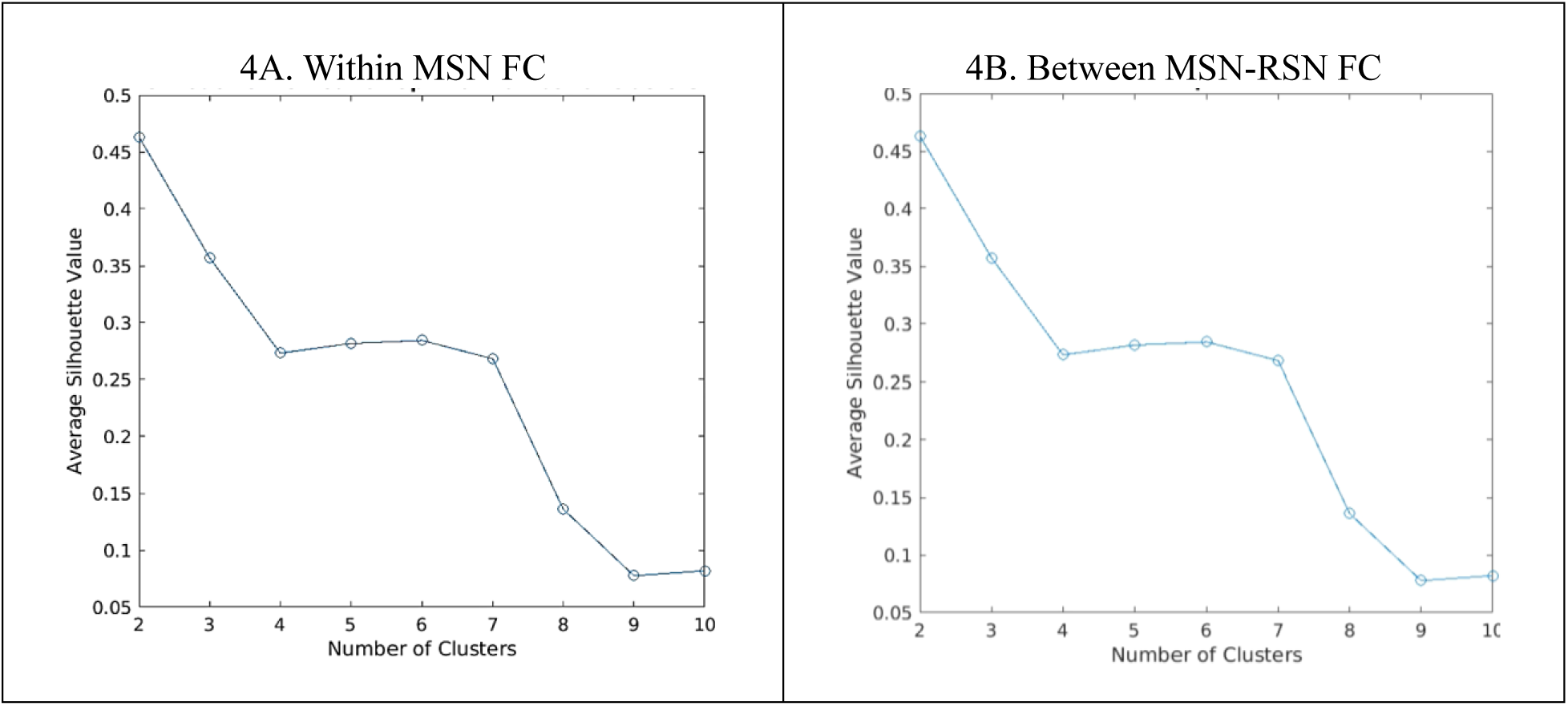
The optimal numbers of clusters in the directed FC within-MSN (4A) and between MSN-RSNs (4B) from silhouette method

## Reference

1 Senkowski D, Engel AK. Multi-timescale neural dynamics for multisensory integration. Nature Reviews Neuroscience 2024 25:9 2024; 25: 625–642.

2 Bizley JK, Jones GP, Town SM. Where are multisensory signals combined for perceptual decision-making? Curr Opin Neurobiol 2016; 40: 31–37.

3 Schroeder CE, Foxe J. Multisensory contributions to low-level, ‘unisensory’ processing. Curr Opin Neurobiol 2005; 15: 454–458.

4 Jones LA, Hills PJ, Dick KM, Jones SP, Bright P. Cognitive mechanisms associated with auditory sensory gating. Brain Cogn 2016; 102: 33–45.

5 Quak M, London RE, Talsma D, Majerus S, Berryhill M, Morey CC et al. A multisensory perspective of working memory. Front Hum Neurosci 2015; 9: 197.

6 Schulz SE, Luszawski M, Hannah KE, Stevenson RA. Sensory Gating in Neurodevelopmental Disorders: A Scoping Review. Res Child Adolesc Psychopathol 2023; 51: 1005–1019.

7 Denervaud S, Gentaz E, Matusz PJ, Murray MM. Multisensory Gains in Simple Detection Predict Global Cognition in Schoolchildren. Scientific Reports 2020 10:1 2020; 10: 1–11.

8 Mason GM, Goldstein MH, Schwade JA. The role of multisensory development in early language learning. J Exp Child Psychol 2019; 183: 48–64.

9 Dionne-Dostie E, Paquette N, Lassonde M, Gallagher A. Multisensory Integration and Child Neurodevelopment. Brain Sci 2015; 5: 32.

10 Stevenson RA, Segers M, Ferber S, Barense MD, Wallace MT. The impact of multisensory integration deficits on speech perception in children with autism spectrum disorders. Front Psychol 2014; 5. doi:10.3389/FPSYG.2014.00379.

11 Isaacs D, Key AP, Cascio CJ, Conley AC, Riordan H, Walker HC et al. Cross-disorder comparison of sensory over-responsivity in chronic tic disorders and obsessive-compulsive disorder. Compr Psychiatry 2022; 113. doi:10.1016/J.COMPPSYCH.2021.152291.

12 Simon DM, Wallace MT. Dysfunction of sensory oscillations in Autism Spectrum Disorder. Neurosci Biobehav Rev 2016; 68: 848–861.

13 Baum SH, Stevenson RA, Wallace MT. Behavioral, perceptual, and neural alterations in sensory and multisensory function in autism spectrum disorder. Prog Neurobiol 2015; 134: 140–160.

14 American Psychiatric Association. Diagnostic and Statistical Manual of Mental Disorders: DSM-V. American Psychiatric Association: Washington DC, 2013.

15 Leekam SR, Nieto C, Libby SJ, Wing L, Gould J. Describing the sensory abnormalities of children and adults with autism. J Autism Dev Disord 2007; 37: 894–910.

16 Tomchek SD, Dunn W. Sensory Processing in Children With and Without Autism: A Comparative Study Using the Short Sensory Profile. The American Journal of Occupational Therapy 2007; 61: 190–200.

17 Robertson CE, Baron-Cohen S. Sensory perception in autism. Nature Reviews Neuroscience 2017 18:11 2017; 18: 671–684.

18 Boddaert N, Chabane N, Gervais H, Good CD, Bourgeois M, Plumet MH et al. Superior temporal sulcus anatomical abnormalities in childhood autism: a voxel-based morphometry MRI study. Neuroimage 2004; 23: 364–369.

19 Redcay E. The superior temporal sulcus performs a common function for social and speech perception: Implications for the emergence of autism. Neurosci Biobehav Rev 2008; 32: 123–142.

20 Cherkassky VL, Kana RK, Keller TA, Just MA. Functional connectivity in a baseline resting-state network in autism. Neuroreport 2006; 17: 1687–1690.

21 Wallace MT, Stevenson RA. The construct of the multisensory temporal binding window and its dysregulation in developmental disabilities. Neuropsychologia 2014; 64: 105–123.

22 Segers M, Bebko JM, Zapparoli BL, Stevenson RA. A pupillometry study of multisensory social and linguistic processing in autism and typical development. Dev Psychol 2020; 56: 2080–2094.

23 Wallace MT, Woynaroski TG, Stevenson RA. Multisensory integration as a window into orderly and disrupted cognition and communication. Annu Rev Psychol 2020; 71: 193–219.

24 Wallace MT, Stevenson RA. The construct of the multisensory temporal binding window and its dysregulation in developmental disabilities. Neuropsychologia 2014; 64: 105–123.

25 Scheerer NE, Pourtousi A, Yang C, Ding Z, Stojanoski B, Anagnostou E et al. Transdiagnostic Patterns of Sensory Processing in Autism and ADHD. J Autism Dev Disord 2022. doi:10.1007/S10803-022-05798-3.

26 Dunn W, Bennett D. Patterns of Sensory Processing in Children with Attention Deficit Hyperactivity Disorder. 101177/1539449202022001022002; **22**: 4–15.

27 Ghanizadeh A. Visual fields in children with attention-deficit/hyperactivity disorder before and after treatment with stimulants. Acta Ophthalmol 2010; 88: e56–e56.

28 de Wit E, van Dijk P, Hanekamp S, Visser-Bochane MI, Steenbergen B, van der Schans CP et al. Same or Different: The Overlap Between Children With Auditory Processing Disorders and Children With Other Developmental Disorders: A Systematic Review. Ear Hear 2018; 39: 1–19.

29 Ghanizadeh A. Screening signs of auditory processing problem: Does it distinguish attention deficit hyperactivity disorder subtypes in a clinical sample of children? Int J Pediatr Otorhinolaryngol 2009; 73: 81–87.

30 Parush S, Sohmer H, Steinberg A, Kaitz M. Somatosensory function in boys with ADHD and tactile defensiveness. Physiol Behav 2007; 90: 553–558.

31 Schulze M, Aslan B, Farrher E, Grinberg F, Shah N, Schirmer M et al. Network-Based Differences in Top–Down Multisensory Integration between Adult ADHD and Healthy Controls—A Diffusion MRI Study. Brain Sciences 2023, Vol 13, Page 388 2023; 13: 388.

32 Tomyshev A, Lebedeva I, Abdullina E, Kaleda V. Deficient Multisensory Integration with concomitant resting-state connectivity in adult ADHD. European Psychiatry 2022; 65: S332–S332.

33 Hare C, Leung B, Zhai G, Li Y, Stevenson R. A Behavioural Investigation of Multisensory Integration in Youth with Attention-Deficit Hyperactivity Disorder. PsyArXiv Preprints. 2024.https://osf.io/preprints/psyarxiv/qvej7_v1 (accessed 26 Jun2025).

34 Hare C. Multisensory Integration in ADHD: A Behavioral and EEG Investigation in Youth and Adults. Electronic Thesis and Dissertation Repository 2024.https://ir.lib.uwo.ca/etd/10391 (accessed 20 Feb2025).

35 Stevenson RA, VanDerKlok RM, Pisoni DB, James TW. Discrete neural substrates underlie complementary audiovisual speech integration processes. Neuroimage 2011; 55: 1339–1345.

36 McCracken HS, Murphy BA, Ambalavanar U, Glazebrook CM, Yielder PC. Source Localization of Audiovisual Multisensory Neural Generators in Young Adults with Attention-Deficit/Hyperactivity Disorder. Brain Sci 2022; 12: 809.

37 Stevenson RA, Geoghegan ML, James TW. Superadditive BOLD activation in superior temporal sulcus with threshold non-speech objects. Exp Brain Res 2007; 179: 85–95.

38 Stevenson RA, Altieri NA, Kim S, Pisoni DB, James TW. Neural processing of asynchronous audiovisual speech perception. Neuroimage 2010; 49: 3308–3318.

39 Choi I, Lee JY, Lee SH. Bottom-up and top-down modulation of multisensory integration. Curr Opin Neurobiol 2018; 52: 115–122.

40 Seth AK, Barrett AB, Barnett L. Granger Causality Analysis in Neuroscience and Neuroimaging. Journal of Neuroscience 2015; 35: 3293–3297.

41 Barnett L, Seth AK. The MVGC multivariate Granger causality toolbox: A new approach to Granger-causal inference. J Neurosci Methods 2014; 223: 50–68.

42 Mclntosh DN, Miller LJ, Shyu V. Development and validation of the short sensory profile. In: Dunn W (ed). Short sensory profile: User’s manual . Psychological Corporation: San Antonio, 1999, pp 59–73.

43 Vanderwal T, Kelly C, Eilbott J, Mayes L, Castellanos F. Inscapes: A movie paradigm to improve compliance in functional magnetic resonance imaging. Neuroimage 2015; 122: 222–232.

44 Esteban O, Markiewicz CJ, Blair RW, Moodie CA, Isik AI, Erramuzpe A et al. fMRIPrep: a robust preprocessing pipeline for functional MRI. Nature Methods 2018 16:1 2018; 16: 111–116.

45 Scheliga S, Kellermann T, Lampert A, Rolke R, Spehr M, Habel U. Neural correlates of multisensory integration in the human brain: An ALE meta-analysis. Rev Neurosci 2023; 34: 223–245.

46 Stevenson RA, Ghose D, Fister JK, Sarko DK, Altieri NA, Nidiffer AR et al. Identifying and Quantifying Multisensory Integration: A Tutorial Review. Brain Topogr 2014; 27: 707–730.

47 Yeo BTT, Krienen FM, Sepulcre J, Sabuncu MR, Lashkari D, Hollinshead M et al. The organization of the human cerebral cortex estimated by intrinsic functional connectivity. J Neurophysiol 2011; 106: 1125–1165.

48 Johnson WE, Li C, Rabinovic A. Adjusting batch effects in microarray expression data using empirical Bayes methods. Biostatistics 2007; 8: 118–127.

49 Leis AM, McSpadden E, Segaloff HE, Lauring AS, Cheng C, Petrie JG et al. K-medoids clustering of hospital admission characteristics to classify severity of influenza virus infection. Influenza Other Respir Viruses 2023; 17: e13120.

50 Raykov YP, Boukouvalas A, Baig F, Little MA. What to Do When K-Means Clustering Fails: A Simple yet Principled Alternative Algorithm. PLoS One 2016; 11: e0162259.

51 Holm S. A Simple Sequentially Rejective Multiple Test Procedure. Scandinavian Journal of Statistics. 1979; 6: 65–70.

52 Dunn W. The short sensory profile. Psychological Corporation: San Antonio, TX, 1999https://www.scirp.org/reference/referencespapers?referenceid=386851 (accessed 6 Jan2025).

53 Froesel M, Cappe C, Ben Hamed S. A multisensory perspective onto primate pulvinar functions. Neurosci Biobehav Rev 2021; 125: 231–243.

54 Sherman SM. The thalamus is more than just a relay. Curr Opin Neurobiol 2007; 17: 417–422.

55 Leh SE, Chakravarty MM, Ptito A. The Connectivity of the Human Pulvinar: A Diffusion Tensor Imaging Tractography Study. Int J Biomed Imaging 2008; 2008: 789539.

56 Cappe C, Rouiller EM, Barone P. Multisensory anatomical pathways. Hear Res 2009; 258: 28–36.

57 Cappe C, Morel A, Barone P, Rouiller EM. The Thalamocortical Projection Systems in Primate: An Anatomical Support for Multisensory and Sensorimotor Interplay. Cerebral Cortex 2009; 19: 2025–2037.

58 Tripathi V, Batta I, Zamani A, Atad DA, Sheth SKS, Zhang J et al. Default mode network functional connectivity as a transdiagnostic biomarker of cognitive function. Biol Psychiatry Cogn Neurosci Neuroimaging 2025. doi:10.1016/J.BPSC.2024.12.016.

59 Harikumar A, Evans DW, Dougherty CC, Carpenter KLH, Michael AM. A Review of the Default Mode Network in Autism Spectrum Disorders and Attention Deficit Hyperactivity Disorder. Brain Connect 2021; 11: 253.

60 Huang S, Li Y, Zhang W, Zhang B, Liu X, Mo L et al. Multisensory Competition Is Modulated by Sensory Pathway Interactions with Fronto-Sensorimotor and Default-Mode Network Regions. The Journal of Neuroscience 2015; 35: 9064.

61 Lombardo M V., Eyler L, Moore A, Datko M, Barnes CC, Cha D et al. Default mode-visual network hypoconnectivity in an autism subtype with pronounced social visual engagement difficulties. Elife 2019; 8. doi:10.7554/ELIFE.47427.

62 Sripada CS, Kessler D, Angstadt M. Lag in maturation of the brain’s intrinsic functional architecture in attention-deficit/hyperactivity disorder. Proc Natl Acad Sci U S A 2014; 111: 14259–14264.

63 Wang K, Li K, Niu X. Altered Functional Connectivity in a Triple-Network Model in Autism With Co-occurring Attention Deficit Hyperactivity Disorder. Front Psychiatry 2021; 12: 736755.

64 Shimizu VT, Bueno OFA, Miranda MC. Sensory processing abilities of children with ADHD. Braz J Phys Ther 2014; 18: 343.

65 Ghanizadeh A. Sensory Processing Problems in Children with ADHD, a Systematic Review. Psychiatry Investig 2010; 8: 89.

66 Siemann JK, Veenstra-VanderWeele J, Wallace MT. Approaches to Understanding Multisensory Dysfunction in Autism Spectrum Disorder. Autism Res 2020; 13: 1430.

67 Uljarević M, Baranek G, Vivanti G, Hedley D, Hudry K, Lane A. Heterogeneity of sensory features in autism spectrum disorder: Challenges and perspectives for future research. Autism Research 2017; 10: 703–710.

68 Green SA, Hernandez L, Bookheimer SY, Dapretto M. Salience Network Connectivity in Autism Is Related to Brain and Behavioral Markers of Sensory Overresponsivity. J Am Acad Child Adolesc Psychiatry 2016; 55: 618–626.e1.

69 Marhenke R, Acevedo B, Sachse P, Martini M. Individual differences in sensory processing sensitivity amplify effects of post-learning activity for better and for worse. Scientific Reports 2023 13:1 2023; 13: 1–9.

70 Green SA, Hernandez L, Bookheimer SY, Dapretto M. Reduced modulation of thalamocortical connectivity during exposure to sensory stimuli in ASD. Autism Res 2016; 10: 801.

71 Hwang K, Bertolero MA, Liu WB, D’Esposito M. The Human Thalamus Is an Integrative Hub for Functional Brain Networks. The Journal of Neuroscience 2017; 37: 5594.

72 Mease RA, Krieger P, Groh A. Cortical control of adaptation and sensory relay mode in the thalamus. Proc Natl Acad Sci U S A 2014; 111: 6798–6803.

## References

1. Esteban O, Markiewicz CJ, Blair RW, et al. fMRIPrep: a robust preprocessing pipeline for functional MRI. Nature Methods 2018 16:1. 2018;16(1):111–116. doi:10.1038/s41592-018-0235-4

2. Gorgolewski K, Burns CD, Madison C, et al. Nipype: A flexible, lightweight and extensible neuroimaging data processing framework in Python. Front Neuroinform. 2011;5:12318. doi:10.3389/FNINF.2011.00013/BIBTEX

3. Tustison NJ, Avants BB, Cook PA, et al. N4ITK: Improved N3 bias correction. IEEE Trans Med Imaging. 2010;29(6):1310–1320. doi:10.1109/TMI.2010.2046908

4. Dale AM, Fischl B, Sereno MI. Cortical Surface-Based Analysis: I. Segmentation and Surface Reconstruction. Neuroimage. 1999;9(2):179–194. doi:10.1006/NIMG.1998.0395

5. Klein A, Ghosh SS, Bao FS, et al. Mindboggling morphometry of human brains. PLoS Comput Biol. 2017;13(2):e1005350. doi:10.1371/JOURNAL.PCBI.1005350

6. Avants BB, Epstein CL, Grossman M, Gee JC. Symmetric diffeomorphic image registration with cross-correlation: Evaluating automated labeling of elderly and neurodegenerative brain. Med Image Anal. 2008;12(1):26–41. doi:10.1016/J.MEDIA.2007.06.004

7. Fonov V, Evans A, McKinstry R, Almli C, Collins D. Unbiased nonlinear average age-appropriate brain templates from birth to adulthood. Neuroimage. 2009;47:S102. doi:10.1016/S1053-8119(09)70884-5

8. Fonov V, Evans AC, Botteron K, Almli CR, McKinstry RC, Collins DL. Unbiased average age-appropriate atlases for pediatric studies. Neuroimage. 2011;54(1):313–327. doi:10.1016/J.NEUROIMAGE.2010.07.033

9. Zhang Y, Brady M, Smith S. Segmentation of brain MR images through a hidden Markov random field model and the expectation-maximization algorithm. IEEE Trans Med Imaging. 2001;20(1):45–57. doi:10.1109/42.906424

10. Cox RW. AFNI: Software for Analysis and Visualization of Functional Magnetic Resonance Neuroimages. Vol 29.; 1996. Accessed September 9, 2020. https://sscc.nimh.nih.gov/sscc/rwcox/papers/CBM_1996.pdf

11. Jenkinson M, Bannister P, Brady M, Smith S. Improved Optimization for the Robust and Accurate Linear Registration and Motion Correction of Brain Images. Neuroimage. 2002;17(2):825–841. doi:10.1006/NIMG.2002.1132

12. Huntenburg J, Gorgolewski K, Anwander A, Margulies D. Evaluating nonlinear coregistration of BOLD EPI and T1w images. In: 20th Annual Meeting of the Organization for Human Brain Mapping. 2014;1. Accessed June 23, 2025. https://pure.mpg.de/rest/items/item_2327525/component/file_2327523/content

13. Wang S, Peterson DJ, Gatenby JC, Li W, Grabowski TJ, Madhyastha TM. Evaluation of field map and nonlinear registration methods for correction of susceptibility artifacts in diffusion MRI. Front Neuroinform. 2017;11:238590. doi:10.3389/FNINF.2017.00017/BIBTEX

14. Treiber JM, White NS, Steed TC, et al. Characterization and Correction of Geometric Distortions in 814 Diffusion Weighted Images. PLoS One. 2016;11(3):e0152472. doi:10.1371/JOURNAL.PONE.0152472

15. Greve DN, Fischl B. Accurate and robust brain image alignment using boundary-based registration. Neuroimage. 2009;48(1):63–72. doi:10.1016/J.NEUROIMAGE.2009.06.060

16. Power JD, Mitra A, Laumann TO, Snyder AZ, Schlaggar BL, Petersen SE. Methods to detect, characterize, and remove motion artifact in resting state fMRI. Neuroimage. 2014;84:320–341. doi:10.1016/J.NEUROIMAGE.2013.08.048

17. Lindquist MA, Geuter S, Wager TD, Caffo BS. Modular preprocessing pipelines can reintroduce artifacts into fMRI data. Hum Brain Mapp. 2019;40(8):2358–2376. doi:10.1002/HBM.24528;JOURNAL:JOURNAL:10970193;WGROUP:STRING:PUBLICATION

18. Tomchek SD, Huebner RA, Dunn W. Patterns of sensory processing in children with an autism spectrum disorder. Res Autism Spectr Disord. 2014;8(9):1214–1224. doi:10.1016/J.RASD.2014.06.006

19. Mangeot SD, Miller LJ, McIntosh DN, et al. Sensory modulation dysfunction in children with attention-deficit-hyperactivity disorder. Dev Med Child Neurol. 2001;43(6):399–406. doi:10.1111/J.1469-8749.2001.TB00228.X

20. Ahn RR, Miller LJ, Milberger S, McIntosh DN. Prevalence of Parents’ Perceptions of Sensory Processing Disorders Among Kindergarten Children. The American Journal of Occupational Therapy. 2004;58(3):287–293. doi:10.5014/AJOT.58.3.287

21. Scheerer NE, Curcin K, Stojanoski B, et al. Exploring sensory phenotypes in autism spectrum disorder. Mol Autism. 2021;12(1):67. doi:10.1186/S13229-021-00471-5

22. Scheerer NE, Pourtousi A, Yang C, et al. Transdiagnostic Patterns of Sensory Processing in Autism and ADHD. J Autism Dev Disord. Published online 2022. doi:10.1007/S10803-022-05798-3

