## Supplemental for "Functional organization of multisensory integration network in children and youth with neurodevelopmental disorders predicts clinical sensory issues"

Supplementary Table1. Definition of Multisensory Network (MSN) connectivity based on ROI pairings

|  | R STG | L STG | L IFG | R Tha | L Tha |
| --- | --- | --- | --- | --- | --- |
| R STG |  | 1 | 2 | 4 | 7 |
| L STG | 11 |  | 3 | 5 | 8 |
| L IFG | 12 | 15 |  | 6 | 9 |
| R Tha | 13 | 16 | 18 |  | 10 |
| L Tha | 14 | 17 | 19 | 20 |  |

R, right hemisphere; L, left hemisphere; STG, superior temporal gyrus; IFG, inferior frontal gyrus; Tha, thalamus; In undirected FC, FC 1 to 10 represent within-MSN FC across five sub-regions. In directed FC, FC 1 to 10 represent directional influence from row to column and FC 11 to 20 represent directional influence from column to row.

Supplementary Table2. Definition of connectivity between Multisensory Network (MSN) and Resting-state Networks (RSNs) based on ROI pairings

MSN to RSN

|  | FPN | | | | | | | | | | | | | | | | DMN | | | | | | | | | | DAN | | | | | | LIM | | | | VAN | | | | | | | | | | | SOM | | VIS | |
| --- | --- | --- | --- | --- | --- | --- | --- | --- | --- | --- | --- | --- | --- | --- | --- | --- | --- | --- | --- | --- | --- | --- | --- | --- | --- | --- | --- | --- | --- | --- | --- | --- | --- | --- | --- | --- | --- | --- | --- | --- | --- | --- | --- | --- | --- | --- | --- | --- | --- | --- | --- |
|  | 01 | 02 | 03 | 04 | 05 | 06 | 07 | 08 | 09 | 10 | 11 | 12 | 13 | 14 | 15 | 16 | 01 | 02 | 03 | 04 | 05 | 06 | 07 | 08 | 09 | 10 | 01 | 02 | 03 | 04 | 05 | 06 | 01 | 02 | 03 | 04 | 01 | 02 | 03 | 04 | 05 | 06 | 07 | 08 | 09 | 10 | 11 | 01 | 02 | 01 | 02 |
| MSN01 | 1 | 6 | 11 | 16 | 21 | 26 | 31 | 36 | 41 | 46 | 51 | 56 | 61 | 66 | 71 | 76 | 81 | 86 | 91 | 96 | 101 | 106 | 111 | 116 | 121 | 126 | 131 | 136 | 141 | 146 | 151 | 156 | 161 | 166 | 171 | 176 | 181 | 186 | 191 | 196 | 201 | 206 | 211 | 216 | 221 | 226 | 231 | 236 | 241 | 246 | 251 |
| MSN02 | 2 | 7 | 12 | 17 | 22 | 27 | 32 | 37 | 42 | 47 | 52 | 57 | 62 | 67 | 72 | 77 | 82 | 87 | 92 | 97 | 102 | 107 | 112 | 117 | 122 | 127 | 132 | 137 | 142 | 147 | 152 | 157 | 162 | 167 | 172 | 177 | 182 | 187 | 192 | 197 | 202 | 207 | 212 | 217 | 222 | 227 | 232 | 237 | 242 | 247 | 252 |
| MSN03 | 3 | 8 | 13 | 18 | 23 | 28 | 33 | 38 | 43 | 48 | 53 | 58 | 63 | 68 | 73 | 78 | 83 | 88 | 93 | 98 | 103 | 108 | 113 | 118 | 123 | 128 | 133 | 138 | 143 | 148 | 153 | 158 | 163 | 168 | 173 | 178 | 183 | 188 | 193 | 198 | 203 | 208 | 213 | 218 | 223 | 228 | 233 | 238 | 243 | 248 | 253 |
| MSN04 | 4 | 9 | 14 | 19 | 24 | 29 | 34 | 39 | 44 | 49 | 54 | 59 | 64 | 69 | 74 | 79 | 84 | 89 | 94 | 99 | 104 | 109 | 114 | 119 | 124 | 129 | 134 | 139 | 144 | 149 | 154 | 159 | 164 | 169 | 174 | 179 | 184 | 189 | 194 | 199 | 204 | 209 | 214 | 219 | 224 | 229 | 234 | 239 | 244 | 249 | 254 |
| MSN05 | 5 | 10 | 15 | 20 | 25 | 30 | 35 | 40 | 45 | 50 | 55 | 60 | 65 | 70 | 75 | 80 | 85 | 90 | 95 | 100 | 105 | 110 | 115 | 120 | 125 | 130 | 135 | 140 | 145 | 150 | 155 | 160 | 165 | 170 | 175 | 180 | 185 | 190 | 195 | 200 | 205 | 210 | 215 | 220 | 225 | 230 | 235 | 240 | 245 | 250 | 255 |

RSN to MSN

|  | FPN | | | | | | | | | | | | | | | | DMN | | | | | | | | | | DAN | | | | | | LIM | | | | VAN | | | | | | | | | | | SOM | | VIS | |
| --- | --- | --- | --- | --- | --- | --- | --- | --- | --- | --- | --- | --- | --- | --- | --- | --- | --- | --- | --- | --- | --- | --- | --- | --- | --- | --- | --- | --- | --- | --- | --- | --- | --- | --- | --- | --- | --- | --- | --- | --- | --- | --- | --- | --- | --- | --- | --- | --- | --- | --- | --- |
|  | 01 | 02 | 03 | 04 | 05 | 06 | 07 | 08 | 09 | 10 | 11 | 12 | 13 | 14 | 15 | 16 | 01 | 02 | 03 | 04 | 05 | 06 | 07 | 08 | 09 | 10 | 01 | 02 | 03 | 04 | 05 | 06 | 01 | 02 | 03 | 04 | 01 | 02 | 03 | 04 | 05 | 06 | 07 | 08 | 09 | 10 | 11 | 01 | 02 | 01 | 02 |
| MSN01 | 256 | 261 | 266 | 271 | 276 | 281 | 286 | 291 | 296 | 301 | 306 | 311 | 316 | 321 | 326 | 331 | 336 | 341 | 346 | 351 | 356 | 361 | 366 | 371 | 376 | 381 | 386 | 391 | 396 | 401 | 406 | 411 | 416 | 421 | 426 | 431 | 436 | 441 | 446 | 451 | 456 | 461 | 466 | 471 | 476 | 481 | 486 | 491 | 496 | 501 | 506 |
| MSN02 | 257 | 262 | 267 | 272 | 277 | 282 | 287 | 292 | 297 | 302 | 307 | 312 | 317 | 322 | 327 | 332 | 337 | 342 | 347 | 352 | 357 | 362 | 367 | 372 | 377 | 382 | 387 | 392 | 397 | 402 | 407 | 412 | 417 | 422 | 427 | 432 | 437 | 442 | 447 | 452 | 457 | 462 | 467 | 472 | 477 | 482 | 487 | 492 | 497 | 502 | 507 |
| MSN03 | 258 | 263 | 268 | 273 | 278 | 283 | 288 | 293 | 298 | 303 | 308 | 313 | 318 | 323 | 328 | 333 | 338 | 343 | 348 | 353 | 358 | 363 | 368 | 373 | 378 | 383 | 388 | 393 | 398 | 403 | 408 | 413 | 418 | 423 | 428 | 433 | 438 | 443 | 448 | 453 | 458 | 463 | 468 | 473 | 478 | 483 | 488 | 493 | 498 | 503 | 508 |
| MSN04 | 259 | 264 | 269 | 274 | 279 | 284 | 289 | 294 | 299 | 304 | 309 | 314 | 319 | 324 | 329 | 334 | 339 | 344 | 349 | 354 | 359 | 364 | 369 | 374 | 379 | 384 | 389 | 394 | 399 | 404 | 409 | 414 | 419 | 424 | 429 | 434 | 439 | 444 | 449 | 454 | 459 | 464 | 469 | 474 | 479 | 484 | 489 | 494 | 499 | 504 | 509 |
| MSN05 | 260 | 265 | 270 | 275 | 280 | 285 | 290 | 295 | 300 | 305 | 310 | 315 | 320 | 325 | 330 | 335 | 340 | 345 | 350 | 355 | 360 | 365 | 370 | 375 | 380 | 385 | 390 | 395 | 400 | 405 | 410 | 415 | 420 | 425 | 430 | 435 | 440 | 445 | 450 | 455 | 460 | 465 | 470 | 475 | 480 | 485 | 490 | 495 | 500 | 505 | 510 |

Fronto-Parietal Network (FPN): FPN01, Left cingulate cortex (L CC); FPN02, Left orbitofrontal cortex (L OFC); FPN03, Left parietal cortex (L PC); FPN04, Left precuneus (L PCu); FPN05, Left dorsal prefrontal cortex (L dPFC); FPN06, Left lateral prefrontal cortex (L lPFC); FPN07, Left medial prefrontal cortex (L mPFC); FPN08, Left ventral prefrontal cortex (L vPFC); FPN09, Left temporal cortex (L TC); FPN10, Right cingulate cortex (R CC); FPN11, Right parietal cortex (R PC); FPN12, Right precuneus (R PCu); FPN13, Right lateral prefrontal cortex (R lPFC); FPN14, Right medial prefrontal cortex (R mPFC); FPN15, Right ventral prefrontal cortex (R vPFC); FPN16, Right temporal cortex (R TC)

Default Mode Network (DMN): DMN01, Left parietal cortex (L PC); DMN02, Left precuneus/posterior cingulate cortex (L PCu/PCC); DMN03, Left prefrontal cortex (L PFC); DMN04, Left parahippocampal cortex (L PHC); DMN05, Left temporal cortex (L TC); DMN06, Right parietal cortex (R PC); DMN07, Right precuneus/posterior cingulate cortex (R PCu/PCC); DMN08, Right dorsal & medial prefrontal cortex (R dmPFC); DMN09, Right ventral prefrontal cortex (R vPFC); DMN10, Right temporal cortex (R TC)

Dorsal Attention Network (DAN): DAN01, Left frontal eye field (L FEF); DAN02, Left posterior cortex (L PC); DAN03, Left precentral cortex (L PreC); DAN04, Right frontal eye field (R FEF); DAN05, Right posterior cortex (R PC); DAN06, Right precentral cortex (R PreC)

Limbic Network (LIM): LIM01, Left orbitofrontal cortex (L OFC); LIM02, Left temporal pole (L TP); LIM03, Right orbitofrontal cortex (R OFC); LIM04, Right temporal pole (R TP)

Ventral Attention Network (VAN): VAN01, Left frontal operculum/insula (L FO/Ins); VAN02, Left medial cortex (L MC); VAN03, Left parietal operculum (L PO); VAN04, Left lateral prefrontal cortex (L lPFC); VAN05, Left temporo-occipital cortex (L TOC); VAN06, Right frontal operculum/insula (R FO/Ins); VAN07, Right medial cortex (R MC); VAN08, Right lateral prefrontal cortex (R lPFC); VAN09, Right ventral prefrontal cortex (R vPFC); VAN10, Right precentral cortex (R PreC); VAN11, Right temporo-occipito-parietal cortex (R TOPC)

Somatomotor Network (SOM): SOM01, Left somatomotor cortex (L SMC); SOM02, Right somatomotor cortex (R SMC)

Visual Network (VIS): VIS01, Left visual cortex (L VC); VIS02, Right visual cortex (R VC)

In undirected FC, FC 1 to 255 represent the influences between MSN and RSNs. In directed FC, FC 1 to 255 represent directional influences from MSN to RSN, while FC 256 to 510 represent directional influences from RSN to MSN.

Supplementary Table 3. Diagnostic differences in the undirected FC within-MSN while controlling for age and sex

| Within-MSN | F | *p*_corrected_ | Partial 𝜂_𝑝_^2^ | Post-hoc |
| --- | --- | --- | --- | --- |
| FC1 (R STG – L STG) | 1.67 | 0.18 | 0.01 |  |
| FC2 (R STG- L IFG) | 0.80 | 0.46 | 0.004 |  |
| FC3 (L STG- L IFG) | 0.43 | 0.64 | 0.002 |  |
| FC4 (R STG – R Tha) | 1.14 | 0.32 | 0.01 |  |
| FC5 (L STG – R Tha) | 3.10 | 0.05* | 0.01 | ADHD>TD |
| FC6 (L IFG – R Tha) | 1.36 | 0.26 | 0.01 |  |
| FC7 (R STG – L Tha) | 1.73 | 0.17 | 0.01 |  |
| FC8 (L STG – L Tha) | 2.95 | 0.06 | 0.01 |  |
| FC9 (L IFG – L Tha) | 0.45 | 0.64 | 0.002 |  |
| FC10 (R Tha – L Tha) | 2.14 | 0.12 | 0.01 |  |

Supplementary Table 4. Diagnostic differences in the undirected FC between MSN-RSN while controlling for age and sex (only FCs identified as significant at *p*_corrected_<.05 are presented)

| RSNs | MSN | Connectivity # (RSN subregion) | F | *p*_corrected_ | Partial 𝜂_𝑝_^2^ | Post-hoc |
| --- | --- | --- | --- | --- | --- | --- |
| FPN | R STG | 1 (L CC) | 3.08 | 0.04 | 0.01 | ADHD>TD |
|  |  | 21 (L dPFC) | 3.38 | 0.04 | 0.02 | ADHD>TD |
|  |  | 46 (R CC) | 3.18 | 0.04 | 0.01 | ADHD>TD |
|  | L STG | 2 (L CC) | 4.69 | 0.01 | 0.02 | ADHD>TD, ASD |
|  |  | 22 (L dPFC) | 4.07 | 0.02 | 0.02 | ADHD>TD |
|  |  | 32 (L mPFC) | 3.05 | 0.05 | 0.01 | ADHD>TD |
|  |  | 42 (L TC) | 5.01 | 0.01 | 0.02 | ADHD>TD |
|  |  | 47 (R CC) | 3.27 | 0.05 | 0.02 | ADHD>TD |
|  |  | 77 (R TC) | 4.05 | 0.02 | 0.02 | ADHD>TD |
|  | L IFG | 3 (L CC) | 6.10 | 0.00 | 0.03 | ADHD>TD |
|  |  | 18 (L PCu) | 3.44 | 0.03 | 0.02 | ADHD>TD |
|  |  | 23 (L dPFC) | 3.24 | 0.04 | 0.01 | ADHD>TD |
|  |  | 43 (L TC) | 3.18 | 0.04 | 0.01 | ADHD>TD |
|  |  | 48 (R CC) | 5.66 | 0.00 | 0.03 | ADHD>TD |
|  |  | 58 (R PCu) | 4.45 | 0.01 | 0.02 | ASD, ADHD>TD |
|  |  | 78 (R TC) | 3.19 | 0.04 | 0.01 | ADHD>TD |
|  | R Tha | 49 (R CC) | 5.25 | 0.01 | 0.02 | ADHD>TD |
|  |  | 59 (R PCu) | 4.56 | 0.01 | 0.02 | ADHD>TD,ASD |
|  | L Tha | 50 (R CC) | 4.14 | 0.02 | 0.02 | ADHD>TD |
| DMN | R STG | 126 (R TC) | 3.48 | 0.03 | 0.02 | ASD,ADHD>TD |
|  | L STG | 92 (L PFC) | 4.72 | 0.01 | 0.02 | ASD,ADHD>TD |
|  |  | 102 (L TC) | 4.85 | 0.01 | 0.02 | ASD,ADHD>TD |
|  |  | 127 (R TC) | 4.25 | 0.02 | 0.02 | ASD,ADHD,TD |
|  | R Tha | 104 (L TC) | 3.43 | 0.03 | 0.02 | ASD,ADHD>TD |
|  | L Tha | 105 (L TC) | 3.19 | 0.04 | 0.01 | ASD,ADHD>TD |
| DAN | R Tha | 149 (R FEF) | 5.86 | 0.00 | 0.03 | ADHD>TD,ASD |
| VAN | R STG | 186 (L PO) | 3.36 | 0.04 | 0.02 | ADHD>TD |
|  |  | 191 (L IPFC) | 3.16 | 0.04 | 0.01 | ADHD>TD |
|  |  | 226 (R PreC) | 3.42 | 0.04 | 0.02 | TD>ASD |
|  | L STG | 187 (L MC) | 3.20 | 0.04 | 0.01 | ADHD>TD |
|  | R Tha | 189 (L MC) | 3.26 | 0.04 | 0.02 | ADHD>TD |
|  |  | 194 (L PO) | 3.08 | 0.05 | 0.01 | ASD,ADHD>TD |
|  |  | 209 (R FO/Ins) | 3.03 | 0.05 | 0.01 | ASD,ADHD>TD |
| SOM | R Tha | 239 (L SMC) | 3.51 | 0.03 | 0.02 | ADHD>TD |
|  |  | 244 (R SMC) | 4.17 | 0.02 | 0.02 | ADHD>TD |
|  | L Tha | 240 (L SMC) | 3.52 | 0.03 | 0.02 | ADHD>TD |
|  |  | 245 (R SMC) | 3.41 | 0.03 | 0.02 | ADHD>TD |

Supplementary Table 5. Associations between sensory processing abilities and the undirected FC within-MSN while controlling for age and sex

| Within-MSN | β | SE | *t* | *p*_corrected_ | Partial *R*^2^ |
| --- | --- | --- | --- | --- | --- |
| FC1 (R STG – L STG) | -0.0005 | 0.001 | -0.88 | 0.38 | 0.00 |
| FC2 (R STG- L IFG) | -0.001 | 0.0005 | -1.43 | 0.16 | 0.01 |
| FC3 (L STG- L IFG) | -0.001 | 0.001 | -1.17 | 0.24 | 0.00 |
| FC4 (R STG – R Tha) | -0.001 | 0.001 | -2.71 | 0.01** | 0.03 |
| FC5 (L STG – R Tha) | -0.001 | 0.0005 | -2.87 | 0.003** | 0.03 |
| FC6 (L IFG – R Tha) | -0.001 | 0.0004 | -3.05 | 0.004** | 0.03 |
| FC7 (R STG – L Tha) | -0.001 | 0.0005 | -2.97 | 0.003** | 0.03 |
| FC8 (L STG – L Tha) | -0.002 | 0.0005 | -3.61 | 0.0002*** | 0.05 |
| FC9 (L IFG – L Tha) | -0.001 | 0.0004 | -2.33 | 0.02* | 0.03 |
| FC10 (R Tha – L Tha) | -0.001 | 0.001 | -1.75 | 0.08 | 0.06 |

Supplementary Table 6. Associations between sensory processing abilities and the undirected FC within-MSN while controlling for age and sex (only FCs identified as significant at *p*_corrected_ <.05 are presented)

| RSN | MSN | Connectivity #  (RSN subregion) | β | SE | *p*_corrected_ | Partial *R*^2^ |
| --- | --- | --- | --- | --- | --- | --- |
| FPN | R STG | 1 (L CC) | -0.001 | 0.0005 | 0.04 | 0.02 |
|  |  | 21 (L dPFC) | -0.001 | 0.0005 | 0.01 | 0.03 |
|  |  | 26 (L lPFC) | -0.001 | 0.0005 | 0.01 | 0.03 |
|  |  | 46 (R CC) | -0.001 | 0.0005 | 0.01 | 0.02 |
|  |  | 56 (R PCu) | -0.001 | 0.0005 | 0.04 | 0.02 |
|  |  | 61 (R lPFC) | -0.001 | 0.0005 | 0.03 | 0.03 |
|  |  | 76 (R TC) | -0.001 | 0.0004 | 0.02 | 0.03 |
|  | L STG | 2 (L CC) | -0.001 | 0.0005 | 0.02 | 0.02 |
|  |  | 12 (L PC) | -0.001 | 0.0005 | 0.03 | 0.03 |
|  |  | 22 (L dPFC) | -0.001 | 0.0005 | 0.01 | 0.02 |
|  |  | 32 (L mPFC) | -0.001 | 0.0005 | 0.01 | 0.03 |
|  |  | 42 (L TC) | -0.001 | 0.0005 | 0.02 | 0.02 |
|  |  | 47 (R CC) | -0.001 | 0.0005 | 0.01 | 0.03 |
|  |  | 62 (R lPFC) | -0.001 | 0.0004 | 0.03 | 0.02 |
|  |  | 77 (R TC) | -0.001 | 0.0004 | 0.003 | 0.04 |
|  | L IFG | 3 (L CC) | -0.002 | 0.0005 | 0.003 | 0.06 |
|  |  | 23 (L dPFC) | -0.001 | 0.0005 | 0.05 | 0.03 |
|  |  | 48 (R CC) | -0.001 | 0.0005 | 0.003 | 0.05 |
|  |  | 58 (R PCu) | -0.001 | 0.0005 | 0.03 | 0.06 |
|  |  | 78 (R TC) | -0.001 | 0.0004 | 0.01 | 0.04 |
|  | R Tha | 4 (L CC) | -0.001 | 0.0004 | 0.02 | 0.04 |
|  |  | 19 (L PCu) | -0.001 | 0.0005 | 0.003 | 0.03 |
|  |  | 34 (L mPFC) | -0.001 | 0.0004 | 0.02 | 0.01 |
|  |  | 39 (L vPFC) | -0.001 | 0.0005 | 0.02 | 0.02 |
|  |  | 44 (L TC) | -0.001 | 0.0004 | 0.01 | 0.02 |
|  |  | 79 (R TC) | -0.001 | 0.0004 | 0.002 | 0.03 |
|  | L Tha | 5 (L CC) | -0.001 | 0.0004 | 0.01 | 0.05 |
|  |  | 20 (L PCu) | -0.001 | 0.0005 | 0.01 | 0.02 |
|  |  | 25 (L dPFC) | -0.001 | 0.0004 | 0.04 | 0.02 |
|  |  | 30 (L lPFC) | -0.001 | 0.0004 | 0.02 | 0.03 |
|  |  | 35 (L mPFC) | -0.001 | 0.0004 | 0.03 | 0.02 |
|  |  | 40 (L vPFC) | -0.001 | 0.0005 | 0.03 | 0.03 |
|  |  | 45 (L TC) | -0.001 | 0.0004 | 0.002 | 0.03 |
|  |  | 50 (R CC) | -0.001 | 0.0004 | 0.01 | 0.04 |
|  |  | 80 (R TC) | -0.001 | 0.0004 | 0.001 | 0.04 |
| DMN | R STG | 81 (L PC) | -0.001 | 0.0005 | 0.02 | 0.03 |
|  |  | 96 (L PHC) | -0.001 | 0.0005 | 0.03 | 0.02 |
|  |  | 101 (L TC) | -0.001 | 0.0005 | 0.01 | 0.05 |
|  |  | 106 (R PC) | -0.001 | 0.0005 | 0.05 | 0.03 |
|  |  | 111 (R PCu/PCC) | -0.001 | 0.0005 | 0.05 | 0.02 |
|  |  | 121 (R vPFC) | -0.001 | 0.0005 | 0.03 | 0.01 |
|  |  | 126 (R TC) | -0.002 | 0.0005 | 0.001 | 0.07 |
|  | L STG | 82 (L PC) | -0.001 | 0.0005 | 0.04 | 0.02 |
|  |  | 87 (L PCu/PCC) | -0.001 | 0.0004 | 0.01 | 0.03 |
|  |  | 92 (L PFC) | -0.001 | 0.0005 | 0.01 | 0.02 |
|  |  | 97 (L PHC) | -0.001 | 0.0005 | 0.004 | 0.02 |
|  |  | 102 (L TC) | -0.002 | 0.0005 | <0.0001 | 0.06 |
|  |  | 127 (R TC) | -0.002 | 0.0005 | 0.0004 | 0.06 |
|  | L IFG | 88 (L PCu/PCC) | -0.001 | 0.0004 | 0.01 | 0.03 |
|  |  | 103 (L TC) | -0.001 | 0.0005 | 0.02 | 0.04 |
|  |  | 128 (R TC) | -0.001 | 0.0005 | 0.01 | 0.05 |
|  | R Tha | 84 (L PC) | -0.001 | 0.0005 | 0.04 | 0.02 |
|  |  | 89 (L PCu/PCC) | -0.001 | 0.0005 | 0.04 | 0.02 |
|  |  | 94 (L PFC) | -0.001 | 0.0005 | 0.02 | 0.02 |
|  |  | 99 (L PHC) | -0.001 | 0.0005 | 0.04 | 0.03 |
|  |  | 104 (L TC) | -0.001 | 0.0004 | 0.003 | 0.04 |
|  |  | 129 (R TC) | -0.001 | 0.0004 | 0.002 | 0.05 |
|  | L Tha | 85 (L PC) | -0.001 | 0.0004 | 0.02 | 0.02 |
|  |  | 90 (L PCu/PCC) | -0.001 | 0.0005 | 0.005 | 0.04 |
|  |  | 95 (L PFC) | -0.001 | 0.0004 | 0.03 | 0.02 |
|  |  | 100 (L PHC) | -0.002 | 0.0005 | 0.0004 | 0.04 |
|  |  | 105 (L TC) | -0.001 | 0.0004 | 0.003 | 0.06 |
|  |  | 110 (R PC) | -0.001 | 0.0004 | 0.04 | 0.02 |
|  |  | 125 (R vPFC) | -0.001 | 0.0005 | 0.01 | 0.03 |
|  |  | 130 (R TC) | -0.001 | 0.0004 | 0.002 | 0.06 |
| DAN | R STG | 141 (L PreC) | -0.001 | 0.0005 | 0.03 | 0.02 |
|  |  | 150 (R FEF) | -0.001 | 0.0004 | 0.02 | 0.02 |
|  |  | 156 (R PreC) | -0.001 | 0.0005 | 0.03 | 0.02 |
|  | L STG | 142 (L PreC) | -0.001 | 0.0005 | 0.04 | 0.02 |
|  |  | 157 (R PreC) | -0.001 | 0.0005 | 0.02 | 0.03 |
|  | L IFG | 133 (L FEF) | -0.001 | 0.0005 | 0.004 | 0.02 |
|  |  | 148 (R FEF) | -0.001 | 0.0005 | 0.04 | 0.02 |
|  | R Tha | 134 (L FEF) | -0.001 | 0.0004 | 0.03 | 0.01 |
|  |  | 144 (L PreC) | -0.001 | 0.0005 | 0.001 | 0.03 |
|  |  | 159 (R PreC) | -0.001 | 0.0005 | 0.03 | 0.01 |
|  | L Tha | 135 (L FEF) | -0.001 | 0.0005 | 0.02 | 0.02 |
|  |  | 145 (L PreC) | -0.001 | 0.0004 | 0.03 | 0.03 |
|  |  | 160 (R PreC) | -0.001 | 0.0004 | 0.03 | 0.02 |
| LIM | L Tha | 165 (L OFC) | -0.001 | 0.0004 | 0.03 | 0.02 |
| VAN | R STG | 206 (R FO/Ins) | -0.001 | 0.0006 | 0.03 | 0.03 |
|  | L STG | 187 (L MC) | -0.001 | 0.0005 | 0.01 | 0.03 |
|  |  | 197 (L lPFC) | -0.001 | 0.0005 | 0.01 | 0.04 |
|  |  | 207 (R FO/Ins) | -0.001 | 0.0005 | 0.04 | 0.03 |
|  |  | 212 (R MC) | -0.001 | 0.0005 | 0.03 | 0.03 |
|  | L IFG | 188 (L MC) | -0.001 | 0.0005 | 0.04 | 0.04 |
|  | R Tha | 184 (L FO/Ins) | -0.001 | 0.0005 | 0.03 | 0.03 |
|  |  | 189 (L MC) | -0.001 | 0.0005 | 0.05 | 0.02 |
|  |  | 194 (L PO) | -0.001 | 0.0004 | 0.01 | 0.03 |
|  |  | 199 (L lPFC) | -0.001 | 0.0005 | 0.02 | 0.02 |
|  |  | 204 (L TOC) | -0.001 | 0.0005 | 0.01 | 0.02 |
|  |  | 209 (R FO/Ins) | -0.001 | 0.0005 | 0.01 | 0.03 |
|  |  | 234 (R TOPC) | -0.001 | 0.0005 | 0.03 | 0.01 |
|  | L Tha | 185 (L FO/Ins) | -0.001 | 0.0005 | 0.02 | 0.04 |
|  |  | 200 (L lPFC) | -0.001 | 0.0005 | 0.03 | 0.03 |
|  |  | 205 (L TOC) | -0.001 | 0.0004 | 0.02 | 0.03 |
|  |  | 210 (F FO/Ins) | -0.002 | 0.0005 | 0.002 | 0.05 |
|  |  | 235 (R TOPC) | -0.001 | 0.0005 | 0.02 | 0.03 |
| SOM | L STG | 237 (L SMC) | -0.001 | 0.0005 | 0.04 | 0.02 |
|  | R Tha | 239 (L SMC) | -0.001 | 0.0005 | 0.02 | 0.03 |
|  |  | 244 (R SMC) | -0.001 | 0.0005 | 0.02 | 0.03 |
|  | L Tha | 240 (L SMC) | -0.001 | 0.0005 | 0.01 | 0.03 |
|  |  | 245 (R SMC) | -0.001 | 0.0005 | 0.005 | 0.02 |
| VIS | R STG | 246 (L VC) | -0.001 | 0.0005 | 0.04 | 0.03 |
|  | L STG | 247 (L VC) | -0.001 | 0.0005 | 0.04 | 0.04 |
|  | R Tha | 249 (L VC) | -0.001 | 0.0005 | 0.02 | 0.01 |
|  | L Tha | 250 (L VC) | -0.001 | 0.0005 | 0.03 | 0.02 |

Supplementary Table 7. Diagnostic differences in the directed FC within-MSN while controlling for age and sex

| Within-MSN directed FC | F | *p*_corrected_ | Partial 𝜂𝑝2 |
| --- | --- | --- | --- |
| FC1 (R STG → L STG) | 0.07 | 0.93 | 0.0003 |
| FC2 (R STG → L IFG) | 0.95 | 0.39 | 0.005 |
| FC3 (L STG → L IFG) | 0.76 | 0.45 | 0.004 |
| FC4 (R STG → R Tha) | 0.07 | 0.94 | 0.0003 |
| FC5 (L STG → R Tha) | 2.19 | 0.11 | 0.01 |
| FC6 (L IFG → R Tha) | 1.15 | 0.31 | 0.006 |
| FC7 (R STG → L Tha) | 1.42 | 0.25 | 0.007 |
| FC8 (L STG → L Tha) | 1.07 | 0.34 | 0.005 |
| FC9 (L IFG → L Tha) | 0.17 | 0.84 | 0.0008 |
| FC10 (R Tha → L Tha) | 1.30 | 0.27 | 0.006 |
| FC11 (L STG → R STG) | 0.22 | 0.81 | 0.001 |
| FC12 (L IFG → R STG) | 1.27 | 0.28 | 0.006 |
| FC13 (R Tha → R STG) | 0.09 | 0.92 | 0.0004 |
| FC14 (L Tha → R STG) | 0.97 | 0.37 | 0.005 |
| FC15 (L IFG → L STG) | 1.10 | 0.33 | 0.005 |
| FC16 (R Tha → L STG) | 0.49 | 0.62 | 0.002 |
| FC17 (L Tha → L STG) | 0.47 | 0.63 | 0.002 |
| FC18 (R Tha → L IFG) | 0.09 | 0.91 | 0.0004 |
| FC19 (L Tha → L IFG) | 1.32 | 0.27 | 0.006 |
| FC20 (L Tha → R Tha) | 0.10 | 0.91 | 0.0005 |

Supplementary Table 8. Diagnostic differences in the directed FC between MSN-RSN while controlling for age and sex (only FCs identified as significant at *p*_corrected_ <.05 are presented)

| Directed FC | | | Connectivity #  (RSN subregion) | F | *p*_corrected_ | Partial 𝜂𝑝2 | Post-hoc |
| --- | --- | --- | --- | --- | --- | --- | --- |
| MSN to RSN | R STG | FPN | 31 (L mPFC) | 4.370296 | 0.01 | 0.02 | ADHD>ASD |
|  | L IFG |  | 3 (L CC) | 5.349348 | 0.0052 | 0.02 | TD>ASD, ADHD |
|  |  |  | 28 (L lPFC) | 4.15231 | 0.0102 | 0.02 | TD>ASD |
|  |  |  | 48 (R CC) | 3.051024 | 0.0436 | 0.01 | TD>ASD |
|  | R Tha |  | 9 (L OFC) | 3.011233 | 0.049 | 0.01 | ADHD>ASD |
|  | L Tha |  | 30 (L lPFC) | 3.619022 | 0.0238 | 0.02 | TD>ASD |
|  | L IFG | DMN | 88 (L PCu/PCC) | 4.440819 | 0.0122 | 0.02 | TD, ASD>ADHD |
|  | R Tha |  | 89 (L PCu/PCC) | 4.774696 | 0.0096 | 0.02 | TD>ASD,ADHD |
|  | R Tha | DAN | 159 (R PreC) | 3.857521 | 0.022 | 0.02 | TD>ASD |
|  | R STG | LIM | 161 (L OFC) | 6.862962 | 0.002 | 0.03 | ASD>TD, ADHD |
|  |  |  | 171 (R OFC) | 6.274968 | 0.003 | 0.03 | ASD>TD,ADHD |
| RSN to MSN | FPN | L STG | 292 (L vPFC) | 3.067299 | 0.0456 | 0.01 | TD,ASD>ADHD |
|  |  |  | 297 (L TC) | 3.305908 | 0.0332 | 0.02 | ASD>ADHD |
|  | DMN | L STG | 372 (R dmPFC) | 5.48667 | 0.0042 | 0.03 | ADHD>TD |
|  |  | L IFG | 348 (L PFC) | 3.890643 | 0.0222 | 0.02 | ASD>ADHD |
|  |  |  | 363 (R PC) | 3.005659 | 0.048 | 0.01 | ASD,ADHD>TD |
|  | DAN | R STG | 386 (L FEF) | 3.284116 | 0.0394 | 0.02 | ADHD>TD |
|  |  | R Tha | 405 (R FEF) | 3.013646 | 0.0432 | 0.01 | ADHD>ASD |

Supplementary Table 9. Associations between sensory processing abilities and the directed FC within-MSN while controlling for age and sex

| Within-MSN | β | SE | *t* | *p*_corrected_ | Partial *R*^2^ |
| --- | --- | --- | --- | --- | --- |
| FC1 (R STG → L STG) | -0.00004 | 0.00008 | -0.49 | 0.62 | 0.002 |
| FC2 (R STG → L IFG) | -0.00002 | 0.00005 | -0.28 | 0.80 | 0.002 |
| FC3 (L STG → L IFG) | 0.00013 | 0.00006 | 2.01 | 0.05* | 0.01 |
| FC4 (R STG → R Tha) | -0.00003 | 0.00006 | -0.41 | 0.69 | 0.02 |
| FC5 (L STG → R Tha) | 0.00021 | 0.00007 | 2.83 | 0.005** | 0.02 |
| FC6 (L IFG → R Tha) | -0.00003 | 0.00006 | -0.42 | 0.68 | 0.002 |
| FC7 (R STG → L Tha) | 0.00007 | 0.00007 | 0.95 | 0.34 | 0.007 |
| FC8 (L STG → L Tha) | 0.00008 | 0.00006 | 1.17 | 0.24 | 0.01 |
| FC9 (L IFG → L Tha) | -0.00001 | 0.00007 | -0.15 | 0.88 | 0.02 |
| FC10 (R Tha → L Tha) | 0.00006 | 0.00006 | 0.91 | 0.35 | 0.02 |
| FC11 (L STG → R STG) | -0.00003 | 0.00006 | -0.50 | 0.61 | 0.01 |
| FC12 (L IFG → R STG) | 0.00000 | 0.00004 | -0.09 | 0.94 | 0.02 |
| FC13 (R Tha → R STG) | -0.00003 | 0.00009 | -0.29 | 0.77 | 0.003 |
| FC14 (L Tha → R STG) | -0.00005 | 0.00010 | -0.48 | 0.63 | 0.003 |
| FC15 (L IFG → L STG) | -0.00003 | 0.00010 | -0.31 | 0.76 | 0.004 |
| FC16 (R Tha → L STG) | 0.00003 | 0.00009 | 0.36 | 0.72 | 0.001 |
| FC17 (L Tha → L STG) | -0.00002 | 0.00007 | -0.26 | 0.78 | 0.003 |
| FC18 (R Tha → L IFG) | -0.00012 | 0.00008 | -1.57 | 0.12 | 0.006 |
| FC19 (L Tha → L IFG) | -0.00008 | 0.00007 | -1.08 | 0.29 | 0.008 |
| FC20 (L Tha → R Tha) | -0.00006 | 0.00005 | -1.15 | 0.26 | 0.01 |

Supplementary Table 10. Associations between sensory processing abilities and the directed FC between MSN-RSN while controlling for age and sex (only FCs identified as significant at *p*_corrected_<.05 are presented)

| Directed FC | | | Connectivity #  (RSN subregion) | β | SE | *t* | *p*_corrected_ | Partial *R*^2^ |
| --- | --- | --- | --- | --- | --- | --- | --- | --- |
| MSN to RSN | L IFG | FPN | 3 (L CC) | 0.003 | 0.001 | 2.60 | 0.01 | 0.02 |
|  |  |  | 48 (R CC) | 0.004 | 0.001 | 2.79 | 0.01 | 0.02 |
|  | R Tha |  | 80 (R TC) | -0.002 | 0.001 | -2.02 | 0.05 | 0.02 |
|  | R Tha | DAN | 159 (R PreC) | 0.008 | 0.003 | 2.34 | 0.02 | 0.01 |
|  | L Tha |  | 135 (L FEF) | -0.004 | 0.002 | -1.99 | 0.05 | 0.01 |
|  |  |  | 150 (R FEF) | -0.003 | 0.002 | -2.01 | 0.04 | 0.01 |
|  |  |  | 155 (R PC) | -0.004 | 0.002 | -2.79 | 0.004 | 0.02 |
|  | L STG | VAN | 222 (R vPFC) | 0.003 | 0.001 | 2.49 | 0.01 | 0.02 |
|  | L IFG |  | 188 (L MC) | 0.003 | 0.001 | 2.04 | 0.04 | 0.01 |
| RSN to MSN | FPN | R STG | 256 (L CC) | -0.005 | 0.002 | -2.13 | 0.03 | 0.02 |
|  |  | R Tha | 275 (L PCu) | 0.007 | 0.003 | 2.51 | 0.01 | 0.02 |
|  | DMN | L STG | 372 (R dmPFC) | -0.008 | 0.002 | -3.43 | 0.0004 | 0.04 |
|  |  | L IFG | 378 (R vPFC) | 0.003 | 0.002 | 2.00 | 0.05 | 0.02 |
|  |  | L Tha | 380 (R vPFC) | 0.004 | 0.002 | 2.22 | 0.03 | 0.02 |
|  | DAN | L Tha | 410 (R PC) | -0.006 | 0.002 | -2.95 | 0.004 | 0.02 |
|  | LIM | R Tha | 429 (R OFC) | -0.004 | 0.002 | -2.30 | 0.02 | 0.02 |

Supplementary Figure 1. Undirected functional connectivity maps within and between MSN and RSNs in three groups.

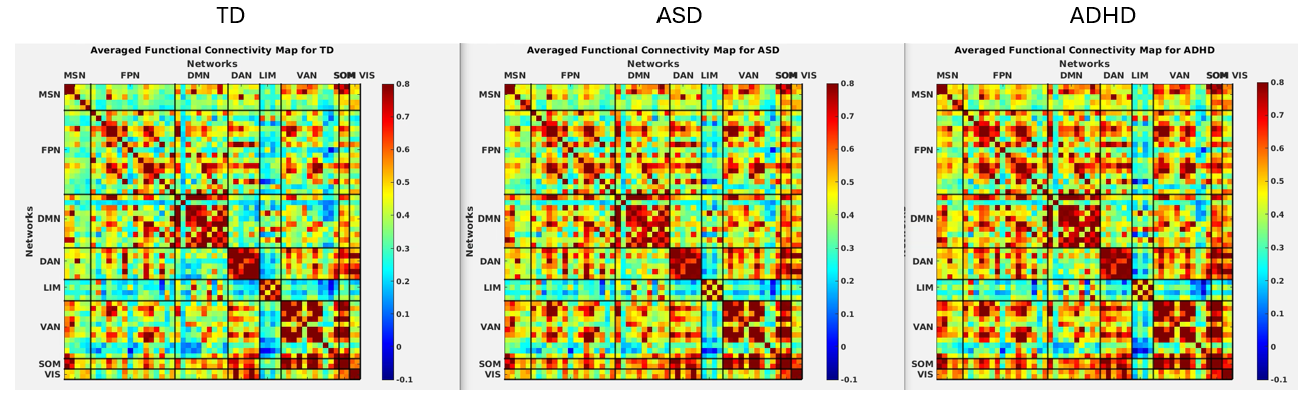

Supplementary Figure 2. The optimal numbers of clusters in the undirected FC within-MSN (2A) and between MSN-RSNs (2B) from elbow method

| 2A. Within MSN FC  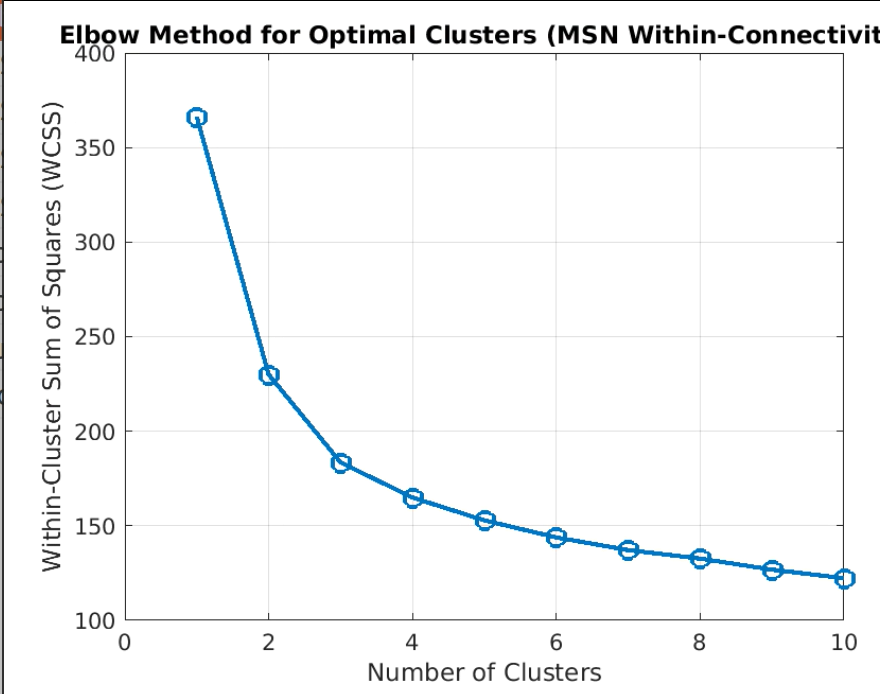 | 2B. Between MSN-RSN FC  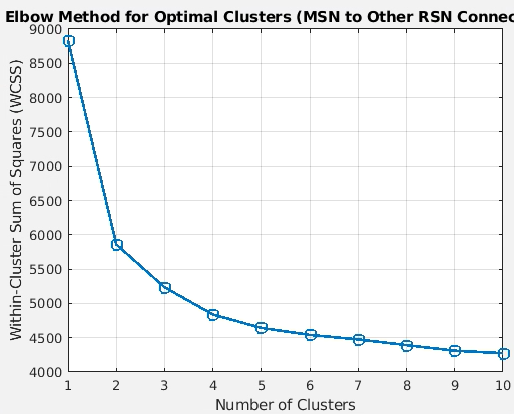 |
| --- | --- |

Supplementary Figure 3. Directed functional connectivity maps within MSN and between MSN and RSNs in three groups.

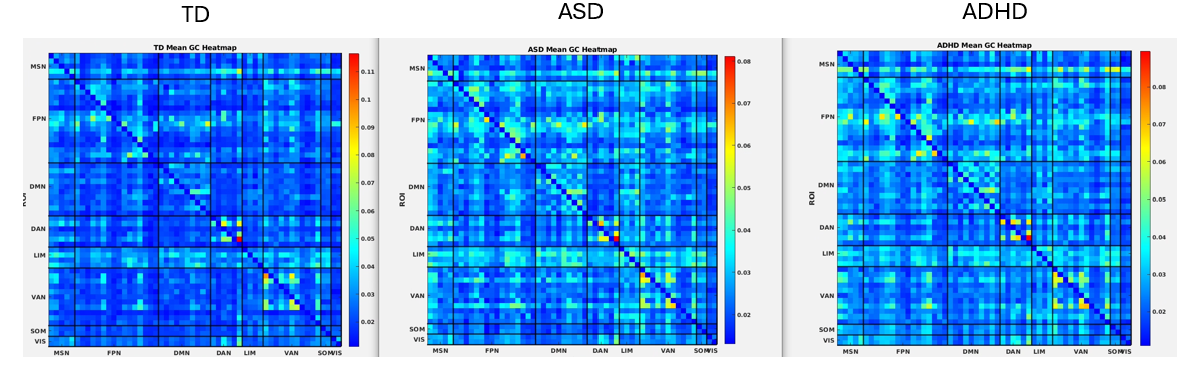

Supplementary Figure 4. The optimal numbers of clusters in the directed FC within-MSN (4A) and between MSN-RSNs (4B) from silhouette method

| 4A. Within MSN FC  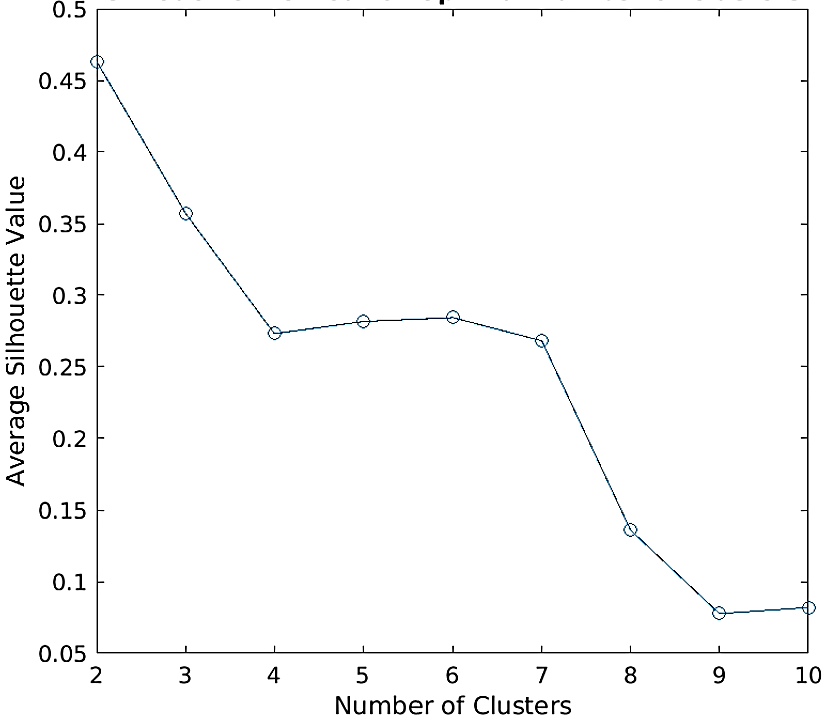 | 4B. Between MSN-RSN FC  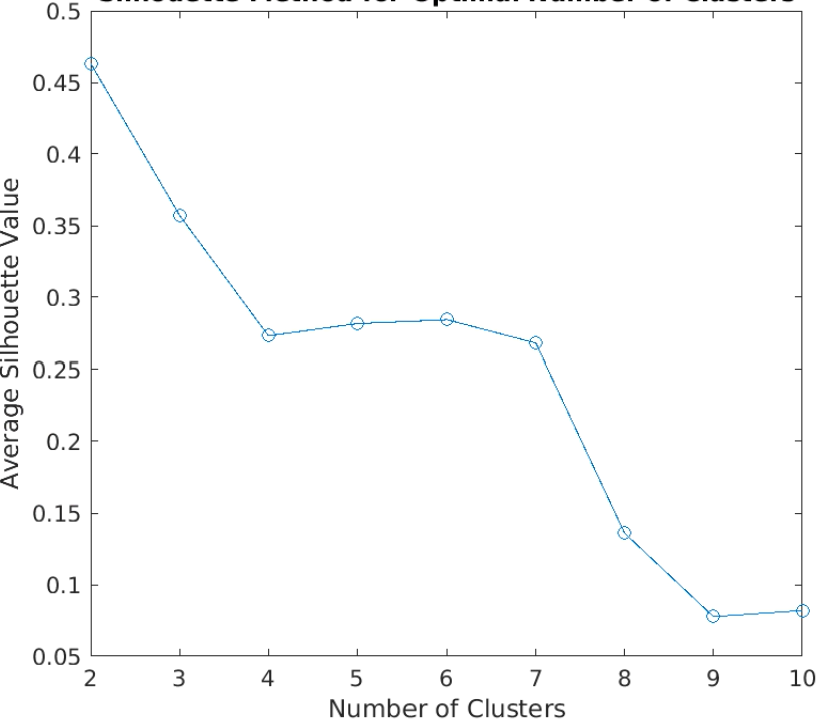 |
| --- | --- |
